# The mammalian cholesterol synthesis enzyme squalene monooxygenase is proteasomally truncated to a constitutively active form

**DOI:** 10.1101/2020.10.12.335414

**Authors:** Hudson W. Coates, Andrew J. Brown

## Abstract

Squalene monooxygenase (SM) is a rate-limiting enzyme of cholesterol synthesis that is oncogenic in a range of cancer types. SM is subject to feedback regulation via cholesterol-induced degradation, which depends on its lipid-sensing N terminal regulatory domain. Here, we characterize an endogenous truncated form of SM and show that it is cholesterol-resistant, and therefore constitutively active. Truncation of SM occurs during its endoplasmic reticulum-associated degradation and requires the proteasome, which partially degrades the SM N-terminus and eliminates cholesterol-sensing elements within this region. Using mutagenesis studies, we demonstrate that partial degradation of SM depends on both an intrinsically disordered region near the truncation site and the stability of the adjacent catalytic domain. Finally, truncation converts SM from an integral to a peripheral ER membrane protein. These findings uncover an additional layer of complexity in the cellular control of cholesterol synthesis and establish SM as the first eukaryotic enzyme known to undergo proteasomal truncation.

## Introduction

Cholesterol is a vital lipid that serves many important functions in mammalian cells, including the maintenance of membrane fluidity and integrity, the assembly of cell surface microdomains for signaling and adhesion, and the synthesis of steroid hormones [1]. Nevertheless, excess cholesterol is cytotoxic and linked with the onset of cardiovascular disease and cancer [2, 3]. It is therefore essential that cells tightly control cholesterol homeostasis by balancing its uptake, synthesis and efflux [4].

The regulation of cholesterol synthesis is especially exquisite, given the energy- and oxygen-intensive nature of the pathway. A critical point at which this regulation is exerted is squalene monooxygenase (SM, also known as squalene epoxidase or SQLE; EC:1.14.14.17), an ER-localized and rate-limiting enzyme responsible for the conversion of squalene to monooxidosqualene [5]. SM is positioned within the branch of the mevalonate pathway that is committed to cholesterol synthesis, contrasting it with the upstream rate-limiting enzyme and well-studied target of the statins, HMG-CoA reductase. Therefore, SM may be an alternative target for the treatment of hypercholesterolemia [6]. Recent years have also seen increasing recognition of SM as oncogenic in a range of malignancies including breast cancer [7], prostate cancer [8] and hepatocellular carcinoma [9]. Moreover, the SM substrate squalene is implicated either as a cytotoxic intermediate [10] or as protective against cancer cell death [11], depending on the cellular context. These reports raise the interesting prospect of targeting SM therapeutically. As the direct pharmacological inhibition of SM is toxic in mammals [12], indirect inhibition by modulating its physiological regulation may be a more viable strategy.

At the transcriptional level, SM expression is controlled by sterol regulatory element-binding proteins, the master regulators of cholesterogenic genes [13, 14]. Acute regulation occurs at the post-translational level, where SM undergoes accelerated degradation in response to increased cholesterol levels [5]. Reciprocally, SM is stabilized by the allosteric binding of squalene [15, 16]. These responses require the N-terminal one hundred amino acids of SM (SM-N100), a regulatory domain that is both necessary and sufficient for cholesterol- and squalene-sensing [5, 15, 17, 18]. The SM-N100 domain is absent from the yeast orthologue of SM, Erg1p, despite high sequence conservation within the SM catalytic domain [5]. This suggests that the lipid-sensing capabilities of SM are unique to higher eukaryotes, in which more nuanced regulation of cholesterol synthesis is required. Cholesterol and squalene affect the stability of SM by modulating its ubiquitination by the E3 ubiquitin ligase membrane-associated RING-CH-type finger 6 (MARCHF6), thereby promoting or preventing its endoplasmic reticulum-associated degradation (ERAD) [15, 19]. Beyond MARCHF6, the ERAD of SM involves additional effectors including the AAA+-type ATPase valosin containing protein (VCP), which extracts client proteins from the ER membrane, the E2 conjugating enzyme Ube2J2, deubiquitinases, and the 26S proteasome [5, 20, 21]. Plasmalogen glycerophospholipids and unsaturated fatty acids also regulate the MARCHF6-mediated degradation of SM [22, 23], implying that SM responds to other classes of lipids. However, further details of the SM ERAD mechanism remain to be elucidated.

Previously, we reported that immunoblotting of SM in HEK293 cell lysates detected full-length SM as well as a lower-molecular weight, putatively truncated form of SM [15]. In the present study, we characterize this SM variant and show that it arises from partial proteasomal degradation of the SM-N100 regulatory domain (referred to herein as proteasomal truncation). This has been described for only two other human proteins, NF-κB and Gli3, where it results in major changes to protein function [24, 25]. In the case of SM, proteasomal truncation depends on an intrinsically disordered region adjacent to the truncation site, as well as the stability of the C-terminal catalytic domain. Truncation yields a constitutively active form of SM that is resistant to cholesterol-accelerated degradation and has an altered ER membrane topology. Therefore, this study uncovers an additional mode by which SM activity is regulated and establishes the first known example of a proteasomally truncated eukaryotic enzyme.

## Results

### A truncated, cholesterol-insensitive form of SM is present in a variety of cell types

We previously reported that anti-SM immunoblotting of HEK293 cell lysates detected full-length SM (∼64 kDa) as well as a lower molecular weight protein (∼55 kDa) that was derived from the *squalene epoxidase* (*SQLE*) gene and strongly stabilized by the SM inhibitor NB-598 [15]. This protein will henceforth be referrezd to as truncated SM (trunSM). Here, we observed expression and NB-598-induced stabilization of trunSM in the commonly used HEK293T and HeLaT cell lines, as well as cell lines derived from tissues that actively synthesize cholesterol: Huh7 (liver), HepG2 (liver) and Be(2)-C (brain) (Fig. 1A). A trunSM-like protein was also detected in the CHO subline CHO-7, where it was stabilized by prolonged NB-598 treatment (Supplementary Fig. S1). These observations confirmed that trunSM production is generalizable to a range of human cell types and the hamster orthologue of SM.

**Figure 1.**
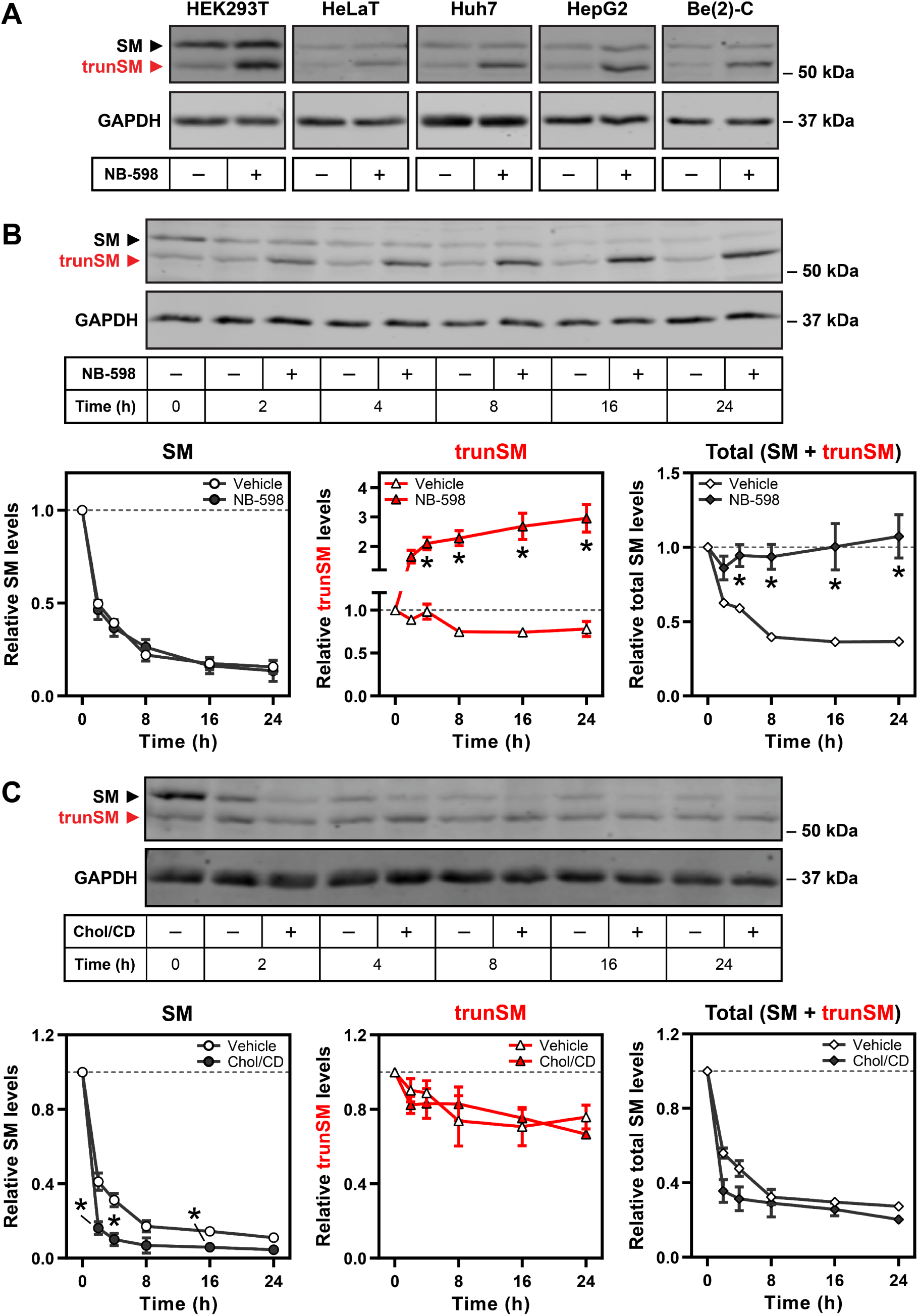
A truncated, cholesterol-insensitive form of SM is present in a variety of cell types. **(A)** The indicated cell lines were treated in the presence or absence of 1 µM NB-598 for 8 h, and immunoblotting was performed for SM and truncated SM (trunSM, red). Immunoblots are representative of *n* ≥ 3 (HEK293T, HeLaT, Huh7, HepG2) or *n* = 1 (Be(2)-C) independent experiments. **(B, C)** HEK293T cells were treated with 10 µg/mL cycloheximide in the presence or absence of (B) µM NB-598 or (C) 20 µg/mL cholesterol-methyl-β-cyclodextrin complexes (Chol/CD) for the indicated times. Graphs depict densitometric quantification of SM levels (left), trunSM levels (center), or total SM levels (SM + trunSM; right) normalized to the 0 h timepoint, which was set to 1 (dotted line). Data presented as mean ± SEM from *n* ≥ 3 independent experiments (*, *p* ≤ 0.05; two-tailed paired *t*-test vs. vehicle condition).

To further characterize trunSM, we examined its stability by treating HEK293T cells with the protein synthesis inhibitor cycloheximide in the presence or absence of NB-598. The trunSM protein was remarkably long-lived: in the absence of NB-598, ∼80% of its starting material remained after 24 h of cycloheximide treatment, compared with only ∼15% of full-length SM (Fig. 1B). NB-598 had no effect on the disappearance of full-length SM but markedly induced trunSM formation, with total SM levels (the sum of full-length SM and trunSM) remaining constant during the treatment. This strongly suggested that trunSM is derived from full-length SM, and that NB-598 promotes this conversion. In a similar experiment, we used co-treatment with cycloheximide and exogenous cholesterol to test if trunSM undergoes the cholesterol-accelerated degradation characteristic of full-length SM [5]. Strikingly, cholesterol had no effect on trunSM levels, whereas accelerated degradation of SM was apparent within 2 h of cholesterol treatment (Fig. 1C). Together, these data indicated that trunSM is induced by NB-598 yet resistant to both basal and cholesterol-induced degradation, raising the possibility that it lacks part or all of the SM-N100 domain. This was consistent with the shift in apparent molecular weight between SM and trunSM, which corresponded to a difference of ∼50–100 amino acids.

### trunSM is not produced by alternative *SQLE* transcripts

The GENCODE- and RefSeq-annotated human genomes each predict a different protein-coding isoform of *SQLE*. These isoforms utilize alternative first exons that substitute the coding sequence of the first 97 amino acids of full-length SM with a two- or 39-amino acid sequence, respectively (Fig. 2A). Given our hypothesis that trunSM lacks the SM-N100 domain, as well as the similarity between the apparent molecular weight of trunSM (∼55 kDa) and the predicted molecular weights of the *SQLE* isoforms (53.1 kDa for the GENCODE isoform, *trunSQLE1*; and 57.6 kDa for the RefSeq isoform, *trunSQLE2*), we sought to confidently rule out the possibility that trunSM arises from alternative *SQLE* transcripts.

**Figure 2.**
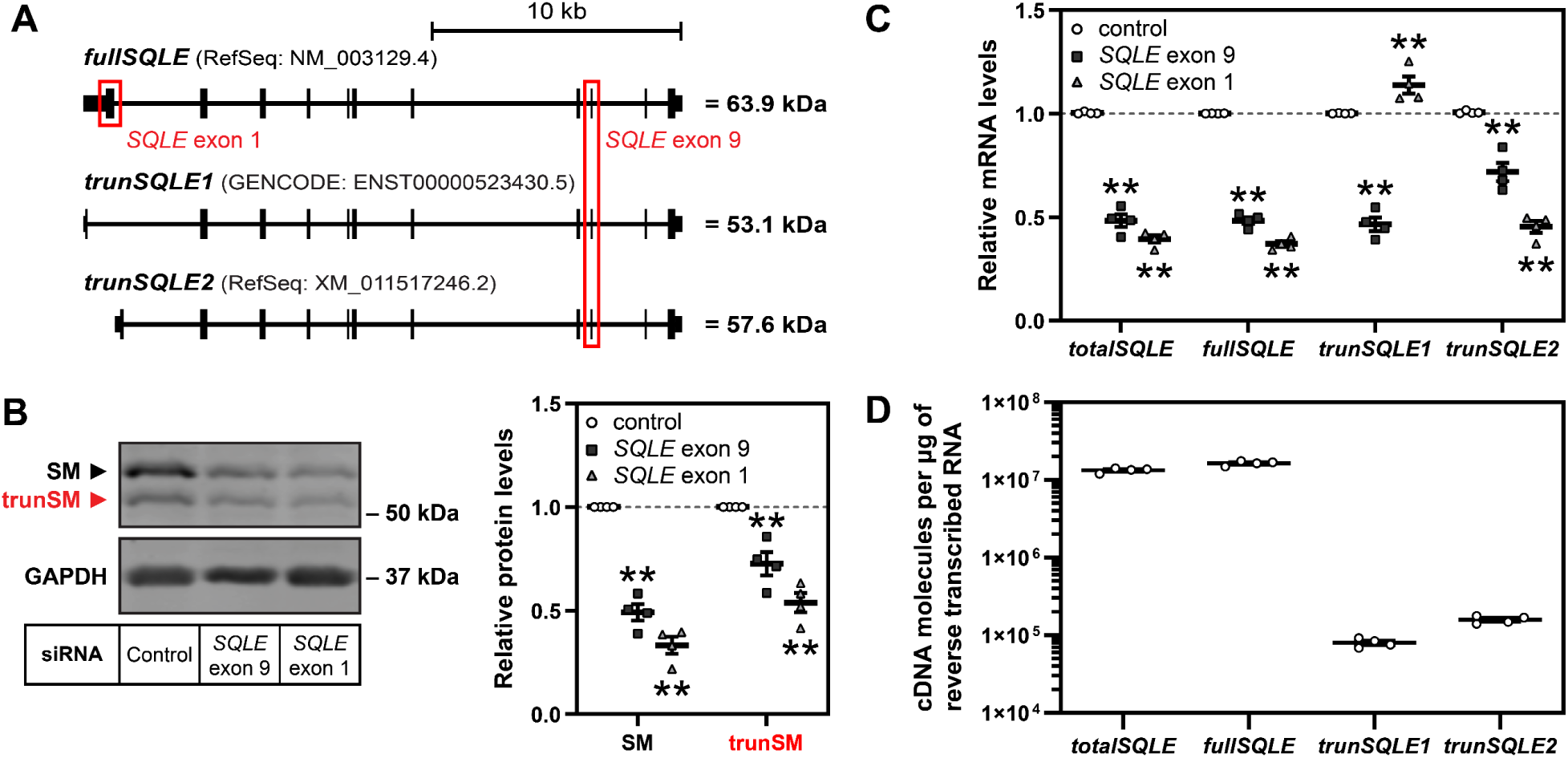
trunSM is not produced by alternative *SQLE* transcripts. **(A)** Schematic of full-length (*fullSQLE*) and alternative protein-coding (*trunSQLE1, trunSQLE2*) *SQLE* transcripts. Exons and untranslated regions are indicated by black bars and siRNA target regions are indicated by red boxes. **(B, C)** HEK293T cells were transfected with the indicated siRNAs for 24 h and refreshed in maintenance medium for a further 24 h. (B) Graph depicts densitometric quantification of SM and trunSM levels normalized to the control siRNA condition, which was set to 1 (dotted line). (C) *SQLE* transcript levels were normalized to *PBGD* housekeeping transcript levels and adjusted relative to the control siRNA condition, which was set to 1 (dotted line). **(D)** Absolute quantification of *SQLE* cDNA levels in control siRNA samples from (C). **(B, C, D)** Data presented as mean ± SEM from *n* = 4 independent experiments, each performed in triplicate for qRT-PCR analysis (**, *p* ≤ 0.01; two-tailed paired *t*-test vs. control siRNA).

To this end, we transfected HEK293T cells with siRNA targeting exon 9 of *SQLE*, which is present in all three *SQLE* isoforms (quantified collectively as *totalSQLE*), or exon 1 of the canonical *SQLE* isoform only (*fullSQLE*; Fig. 2A). Both siRNAs reduced trunSM protein levels (Fig. 2B) whereas *trunSQLE1* mRNA expression was downregulated by only exon 9 siRNA, ruling out this isoform as giving rise to trunSM (Fig. 2C). Unexpectedly, *trunSQLE2* mRNA expression was downregulated by exon 1 siRNA, perhaps due to the presence of the siRNA target sequence in an unannotated 3′-untranslated region of this transcript. To determine the likelihood of *trunSQLE2* accounting for trunSM formation, we next performed absolute quantification of *SQLE* cDNA. Full-length *SQLE* cDNA comprised the great majority of *SQLE* transcripts (∼1.6×10^7^ cDNA copies per µg of reverse transcribed RNA), while *trunSQLE1* and *trunSQLE2* cDNA were less abundant by over two orders of magnitude (∼8.0×10^4^ and ∼1.5×10^5^ cDNA copies, respectively) (Fig. 2D). Given that (1) trunSM protein levels are comparable to full-length SM (Fig. 2B), (2) NB-598-induced accumulation of trunSM occurs in the presence of the protein synthesis inhibitor cycloheximide (Fig. 1B), and (3) we later found that ectopic SM also produces a trunSM-like fragment (Fig. 3A), we concluded that trunSM is highly unlikely to be derived from lowly-abundant *SQLE* isoforms.

**Figure 3.**
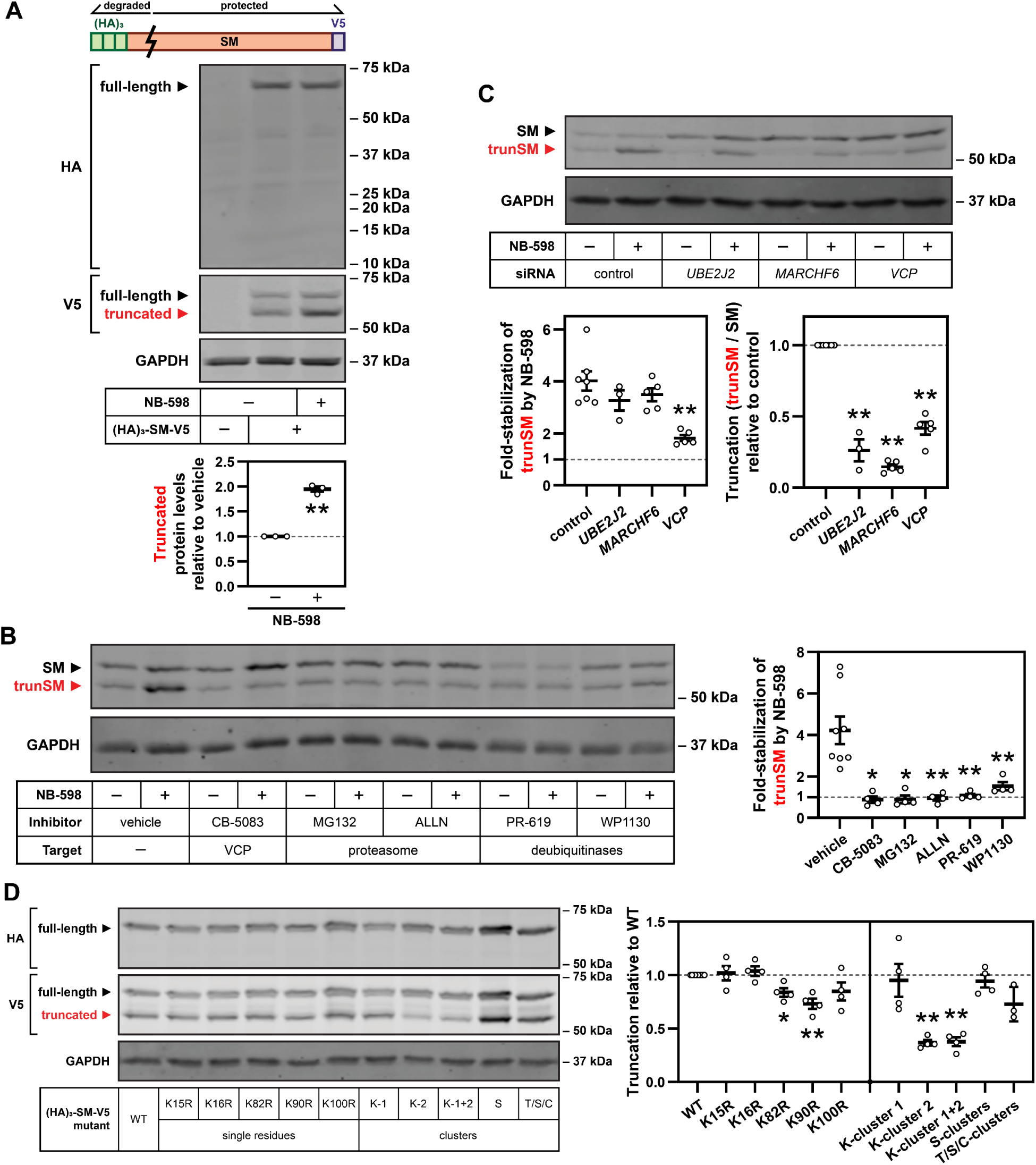
trunSM arises from partial proteasomal degradation of the SM N-terminus. **(A)** HEK293T cells were transfected with empty vector or pCMV-(HA)_3_-SM-V5 expression vector for 24 h, refreshed in maintenance medium for 16 h, and treated in the presence or absence of 1 µM NB-598 for 8 h. Lysates were separated by 4–15% gradient Tris-glycine SDS-PAGE. Graph depicts densitometric quantification of truncated protein levels normalized to the vehicle condition, which was set to 1 (dotted line). **(B)** HEK293T cells were treated with 5 µM CB-5083, 20 µM MG132, 25 µg/mL ALLN, 40 µM PR-619 or 10 µM WP1130, in the presence or absence of 1 µM NB-598, for 8 h. Graph depicts densitometric quantification of trunSM stabilization by NB-598. **(C)** HEK293T cells were transfected with the indicated siRNAs for 24 h, refreshed in maintenance medium for 16 h, and treated in the presence or absence of 1 µM NB-598 for 8 h. Graphs depict densitometric quantification of (left) trunSM stabilization by NB-598, or (right) SM truncation normalized to the control siRNA condition, which was set to 1 (dotted line). **(D)** HEK293T cells were transfected with the indicated constructs for 24 h and refreshed in maintenance medium for 24 h. Graph depicts densitometric quantification of (HA)_3_-SM-V5 truncation normalized to the wild-type (WT) construct, which was set to 1 (dotted line). Cluster mutations: K-cluster 1 (K15R, K16R); K-cluster 2 (K82R, K90R, K100R); K-cluster 1+2 (K15R, K16R, K82R, K90R, K100R); S-clusters (S59A, S61A, S83A, S87A); T/S/C-clusters (T3A, T9A, T11A, S43A, C46A, S59A, S61A, S67A, S71A, S83A, S87A). **(A, B, C, D)** Data presented as mean ± SEM from *n* ≥ 3 independent experiments (*, *p* ≤ 0.05; **, *p* ≤ 0.01, two-tailed paired *t*-test vs. [A, B] vehicle, [C] control siRNA or [D] WT).

### trunSM arises from partial proteasomal degradation of the SM N-terminus

To determine if trunSM is a proteolytic product of full-length SM, we transfected HEK293T cells with SM fused to N-terminal (HA)_3_ and C-terminal V5 epitope tags ([HA]_3_-SM-V5). Immunoblotting detected two C-terminally-tagged proteins with molecular weights corresponding to SM and trunSM, the latter of which was stabilized by NB-598 (Fig. 3A). Only the full-length protein was N-terminally tagged, confirming that the trunSM-like fragment lacks the SM N-terminus. Interestingly, we were unable to recover a low-molecular weight, N-terminally tagged fragment, suggesting that the SM N-terminus undergoes complete proteolysis during truncation. To estimate the truncation site, we inserted a FLAG epitope tag at various positions within the (HA)_3_-SM-V5 construct and monitored for its appearance in the truncated fragment. Truncation eliminated the FLAG tag when it was inserted after SM residue 60 but not residue 70 (Supplementary Fig. S2A), implying that truncation occurs between these two residues. Therefore, trunSM lacks part of the SM-N100 regulatory domain but retains the full C-terminal catalytic domain.

Given that SM truncation does not yield an intact N-terminal fragment (Fig. 3A), we hypothesized that trunSM formation requires the proteasome and, by extension, the ERAD of SM. ERAD effectors involved in the proteasomal degradation of SM include the AAA+-type ATPase VCP, the E3 ubiquitin ligase MARCHF6 and its associated E2 conjugating enzyme Ube2J2, and unidentified deubiquitinases [19, 21]. Treating HEK293T cells with VCP, proteasome or deubiquitinase inhibitors blocked the NB-598-induced accumulation of trunSM (Fig. 3B), confirming that a functional ERAD pathway and the proteasome are required for truncation. To corroborate this finding, we performed siRNA-mediated knockdown of ERAD effectors. Knockdown of *VCP* similarly blunted the NB-598-induced accumulation of trunSM, whilst *UBE2J2* or *MARCHF6* knockdown had no effect (Fig. 3C, left). However, we noted that in the absence of NB-598, all three knockdowns had greatly reduced the basal truncation of SM (Fig. 3C, right; expressed as the ratio between trunSM and full-length SM levels). This suggested that while Ube2J2 and MARCHF6 are the major E2 and E3 proteins required for SM truncation under normal conditions, other proteins can compensate for their absence during NB-598-stimulated truncation. This contrasted with the apparent absolute requirement for VCP. A lysosome-dependent route for SM degradation has been proposed [9]; however, inhibitors of lysosomal acidification had no effect on trunSM formation (Supplementary Fig. S2B), further supporting an ERAD-dependent mechanism.

As ERAD requires substrate ubiquitination, we next sought to identify whether a ubiquitin signal controls truncation. We predicted that this signal would occur within the SM-N100 regulatory domain, given that SM is truncated at its N-terminus. While mutation of Lys-82 and the published ubiquitination site Lys-90 [26] slightly reduced the truncation of the (HA)_3_-SM-V5 construct, a more marked effect was observed upon combined mutation of the Lys-82/90/100 cluster (Fig. 3D; Supplementary Fig. S2C), implying functional redundancy amongst these residues. This reduction in truncation was not compounded by additional mutation of Lys-15/16, residues located nearer to the SM N-terminus, suggesting that Lys-82, Lys-90 and Lys-100 are most critical for truncation. We previously showed that threonine, serine and cysteine residues within SM-N100 contribute to the cholesterol-accelerated degradation of SM, with Ser-83 serving as a non-canonical ubiquitination site [20] (Supplementary Fig. S2C). However, mutating these residues did not affect truncation (Fig. 3D). Given that lysine residues within the SM-N100 domain are not required for cholesterol-accelerated degradation of SM [5, 20], this indicated that the proteasomal truncation of SM depends on a distinct ubiquitin signal.

### SM truncation depends on an intrinsically disordered region and the stability of the catalytic domain

Few other substrates of partial proteasomal degradation are known. However, two features are associated with truncation: (1) a low-complexity sequence [27] or (2) high intrinsic disorder [28]. In both cases, the region must be adjacent to a tightly folded domain that is resistant to proteasomal unfolding and degradation, allowing an opportunity for substrate release [27–29]. To investigate whether these features could account for SM truncation, we analyzed the SM protein sequence using predictors of sequence complexity and intrinsic disorder. Four short regions of low sequence complexity were located throughout SM, including one within the SM-N100 domain (residues 51–62; Fig. 4A). Deletion of this region (Δ50–60) slightly reduced the truncation of the (HA)_3_-SM-V5 construct but did not alter the size of the truncated fragment (Supplementary Fig. S3A), further supporting the idea that truncation occurs after residue 60. More strikingly, we identified a highly intrinsically disordered region between residues 83–120 (Fig. 4A), adjacent to the predicted truncation site. By contrast, residues in the C-terminal direction of this region, comprising the SM catalytic domain, were highly ordered.

**Figure 4.**
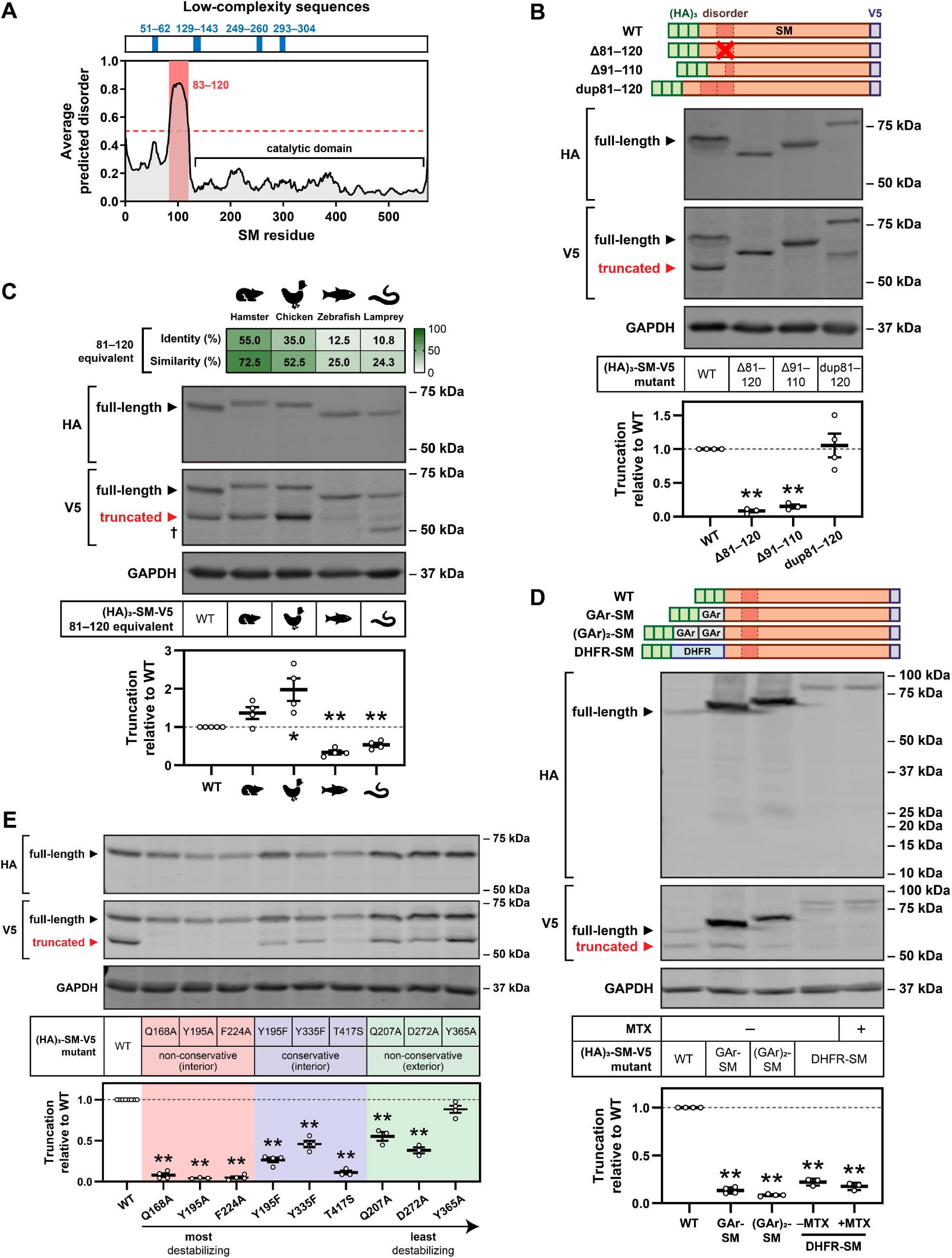
SM truncation depends on an intrinsically disordered region and the stability of the catalytic domain. **(A)** Low-complexity regions (blue) and intrinsically disordered regions (red) within the SM protein sequence. **(B, C, E)** HEK293T cells were transfected with the indicated constructs for 24 h and refreshed in maintenance medium for 24 h. (C) Dagger indicates an additional non-trunSM fragment. **(D)** HEK293T cells were transfected with the indicated constructs for 24 h, refreshed in maintenance medium for 16 h, and treated in the presence or absence of 10 µM methotrexate (MTX) for 8 h. Lysates were separated by 4–15% gradient Tris-glycine SDS-PAGE. **(B – E)** Graphs depict densitometric quantification of (HA)_3_-SM-V5 truncation normalized to the WT construct, which was set to 1 (dotted line). Data presented as mean ± SEM from *n* ≥ 3 independent experiments (*, *p* ≤ 0.05; **, *p* ≤ 0.01, two-tailed paired *t*-test vs. WT).

Supporting the importance of the disordered region in partial degradation of SM, its deletion (Δ81–120) abolished truncation (Fig. 4B). Halving the length of the disordered region (Δ91–110) also prevented truncation, whilst tandem duplication of the disordered region (dup81– 120) had little effect, implying that a minimum length of intrinsic disorder is required for this process. We also noted that the apparent molecular weight of the truncated fragment increased when the disordered region was duplicated, suggesting that the truncation site remained unchanged despite the extended disorder length. As the region corresponding to residues 81–120 is highly disordered in SM orthologues from Chinese hamster, chicken, zebrafish and sea lamprey, despite their differing levels of sequence conservation (Fig. 4C, Supplementary Fig. S3B and S3C), we next tested the effect of substituting these regions into human SM. Truncation was maintained or even enhanced in constructs derived from Chinese hamster and chicken, and approximately halved in constructs derived from zebrafish and sea lamprey SM (Fig. 4C). The persistence of truncation in all four mutant constructs indicated that the intrinsically disordered nature of the 81–120 region is sufficient to promote truncation, although sequence-specific features may have an accessory function.

The proteasome typically engages and initiates degradation from intrinsically disordered regions of its substrates [30]. Therefore, we considered if residues 81–120 of SM are an internal proteasomal engagement site that results in preferential degradation of the N-terminus. A similar mechanism has been reported for other substrates of partial degradation [29]. To test this, we generated N-terminal fusions of SM with two proteins that impede proteasomal processivity: a 30-amino acid glycine-alanine repeat (GAr) from the Epstein-Barr virus nuclear antigen-1 [31], or dihydrofolate reductase (DHFR), which becomes tightly folded and resistant to degradation upon the binding of its ligand methotrexate [32]. We reasoned that if degradation were initiated internally, these fusions would not block truncation but rather protect the N-terminus from complete degradation. However, we found that the fusion of GAr sequences dramatically ablated truncation, and we were unable to recover N-terminal fragments of the expected molecular weight (10–15 kDa; Fig. 4D). Fusion of DHFR similarly reduced truncation, and its further stabilization by methotrexate did not rescue the N-terminus from degradation. This indicated that partial proteasomal degradation of SM is initiated from the N-terminus rather than an internal site. To support this conclusion, we further manipulated the SM N-terminus by sequentially removing HA epitope tags from the (HA)_3_-SM-V5 construct. These tags have a propensity for intrinsic disorder [33] and may enhance proteasomal engagement at the N-terminus. As expected, this led to a stepwise reduction in SM truncation (Supplementary Fig. S3D), confirming that truncation proceeds from the N-terminus.

All known examples of partial proteasomal degradation require a tightly folded domain adjacent to the truncation site. Given that the SM catalytic domain has a compact structure [34] and is predicted to be highly ordered (Fig. 4A), we considered the possibility that its stability is also essential for truncation. Supporting this idea was our earlier observation that NB-598 treatment rapidly accumulates trunSM (Fig. 1B). NB-598 is a potent, tight-binding inhibitor of SM that strongly stabilizes the catalytic domain [34], likely increasing its resistance to proteasomal unfolding. To examine the inverse situation, we generated point mutations within the catalytic domain based on the crystal structure of SM [34]. Our rationale was that non-conservative substitutions in the domain interior would be more destabilizing than conservative substitutions, which would in turn be more destabilizing than mutations on the domain exterior. As expected, most of the substitutions significantly reduced SM truncation, and those that were both internal and non-conservative tended to have a larger effect than those which were conservative or external (Fig. 4E). Of note, the non-conservative mutation of Tyr-195 (Y195A) ablated truncation to a greater extent than its conservative equivalent (Y195F). This strongly suggested that catalytic domain stability is required for partial degradation. Taken together, our data indicate that the truncation of SM depends on two major structural features: the 81–120 disordered region and the stability of the SM catalytic domain.

### trunSM adopts an altered ER membrane topology

The stability and cholesterol-resistance of trunSM (Fig. 1C), as well as the preservation of the entire SM catalytic domain following truncation (Supplementary Fig. S2A), suggested that it would act as a constitutively active form of SM. Previous studies have established that SM lacking the SM-N100 domain retains catalytic activity [34, 35]; therefore, trunSM is highly likely to be active. However, this is contingent on trunSM maintaining the ER localization of full-length SM. Fractionation of HEK293T cell lysates revealed that like full-length SM [5], trunSM is membrane-associated (Fig. 5A). However, a greater proportion of trunSM was found in the cytoplasmic fraction compared with full-length SM, particularly in the absence of NB-598. This suggested that trunSM is more loosely bound to the membrane than full-length SM, possibly due to the loss of the SM-N100 re-entrant loop (residues ∼20–40). To investigate this further, membranes were isolated and treated with aqueous buffer (control), 1% SDS (solubilizing), 0.1 M Na_2_CO_3_ (high pH) or 1 M NaCl (high salt). Solubilizing conditions disrupt the membrane association of all membrane proteins, while high-pH or high-salt conditions release peripheral membrane proteins (in the latter case, those associated *via* electrostatic interactions) [36, 37]. Both full-length SM and trunSM remained membrane-associated under aqueous or high-salt conditions and were released into the supernatant fraction under solubilizing conditions (Fig. 5B). A similar distribution was observed for SM-N100-GFP-V5, a fusion construct that includes the SM-N100 re-entrant loop [18]. However, unlike full-length SM and SM-N100-GFP-V5, the membrane association of trunSM was readily disrupted by high-pH conditions. This suggested that the loss of the SM-N100 re-entrant loop renders trunSM a peripheral ER membrane protein.

**Figure 5.**
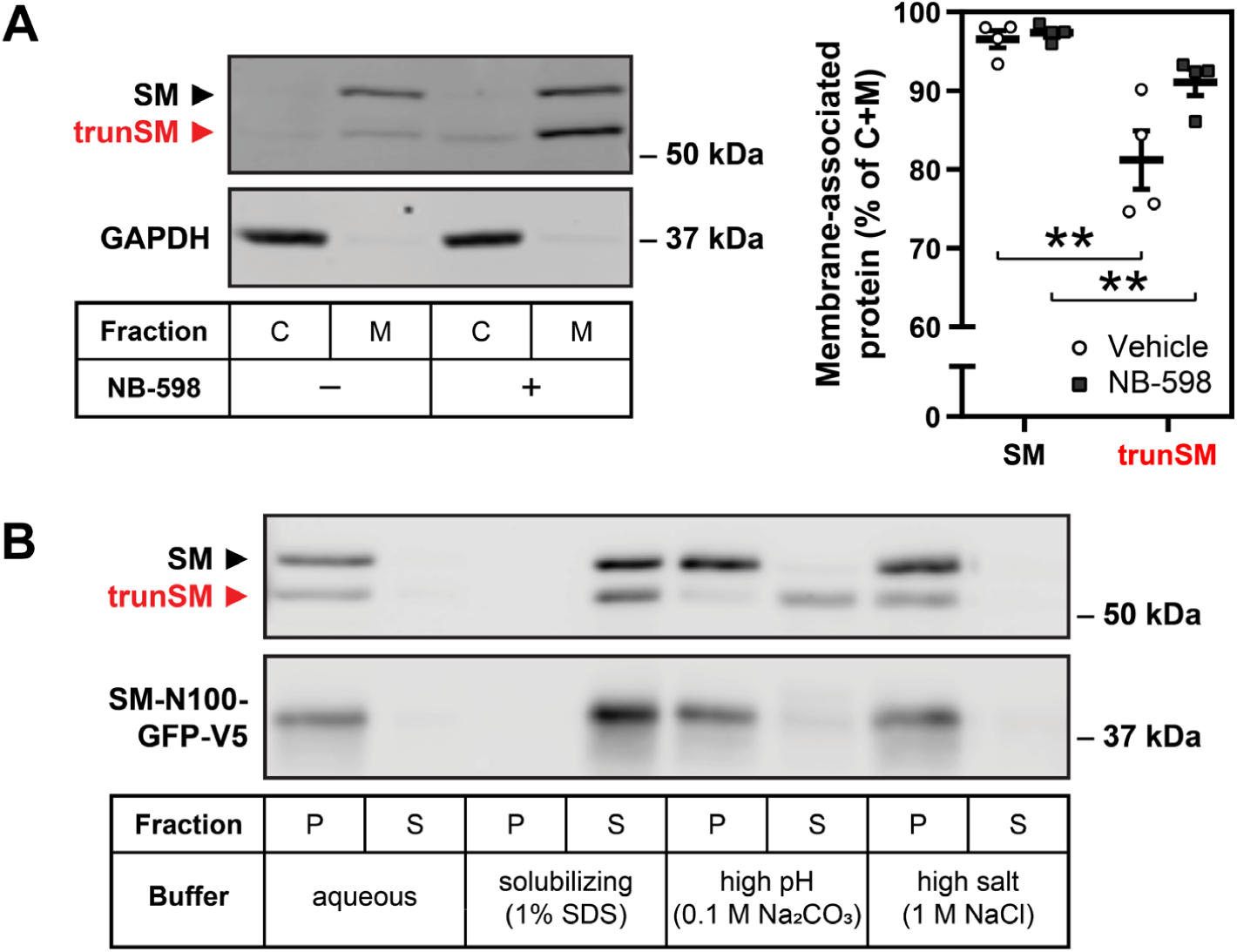
trunSM adopts an altered ER membrane topology. **(A)** HEK293T cells were treated in the presence or absence of 1 µM NB-598 for 8 h, and cytosolic (C) or membrane (M) fractions were isolated. Graph depicts the proportion of overall protein (C + M) found in the membrane fraction. Data presented as mean ± SEM from *n* = 4 independent experiments (**, *p* ≤ 0.01, two-tailed paired *t*-test vs. SM). **(B)** HEK293T cells were transfected with pTK-SM-N100-GFP-V5 for 24 h and refreshed in maintenance medium for a further 24 h. Membrane fractions were isolated and treated as indicated, followed by collection of pellet (P) and supernatant (S) fractions. Immunoblot is representative of *n* = 3 independent experiments.

## Discussion

Feedback regulation of SM protein levels is conferred by its lipid-sensing SM-N100 domain, which contains structural elements required for cholesterol-accelerated degradation. In this study, we characterized a truncated form of SM (trunSM) that is produced by partial proteolysis of the SM-N100 domain. This renders trunSM long-lived, cholesterol-resistant, and, as the SM-N100 domain is not required for catalysis [34, 35], constitutively active. Truncation requires ERAD and the proteasome and depends on two features of SM: intrinsic disorder within the 81–120 region and the stability of the adjacent catalytic domain. Furthermore, the loss of a membrane-embedded region at the N-terminus causes trunSM to adopt a peripheral association with the ER membrane. These findings establish a new mechanism affecting the abundance and activity of SM, with likely consequences for the homeostatic control of cholesterol synthesis.

### Proteasomal truncation of SM

The SM-N100 domain contains two cholesterol-sensing elements that enable its accelerated degradation: a re-entrant loop spanning residues ∼15–40 that undergoes a conformational change in the presence of excess cholesterol [18], and a membrane-associated amphipathic helix from residues 62–73 that deforms and is ejected from the ER membrane under similar conditions [17]. Truncation of the SM N-terminus eliminates the SM-N100 re-entrant loop and likely disrupts the conformation and function of the nearby amphipathic helix, accounting for the longevity and cholesterol-resistance of the trunSM fragment (Fig. 1C). This reinforces the importance of these two structural features for the metabolic regulation of full-length SM, as they have largely been studied only in the context of the isolated SM-N100 domain [17, 18]. Loss of the membrane-embedded re-entrant loop also renders trunSM a peripheral membrane protein (Fig. 5B), bound to the ER membrane *via* two C-terminal helices [34]. Proteomic studies have shown that SM partitions to lipid droplets [38, 39], and it is possible that the peripheral membrane association of trunSM makes it more suited to the lipid droplet monolayer than its full-length counterpart. This possibility warrants further consideration given the predicted constitutive activity of trunSM and the lipid droplet localization of lanosterol synthase, the cholesterol synthesis enzyme immediately downstream of SM [38, 39].

Using pharmacological and genetic approaches we found that the truncation of SM, like its cholesterol-regulated degradation [5], occurs through proteasomal ERAD and requires Ube2J2, MARCHF6 and VCP (Fig. 3B and 3C; Fig. 6). However, the exact mechanism is distinct. Truncation is not stimulated by cholesterol (Fig. 1C), depends on a cluster of lysine residues (Lys-82/90/100) that is dispensable for cholesterol regulation [5, 20], and is independent of atypical cholesterol-dependent ubiquitination sites within SM-N100 (Fig. 3D) [20]. Instead, truncation may occur for a subset of SM molecules undergoing a basal degradation route. This is supported by our finding that upon stabilization of SM by NB-598, complete degradation ceases and all SM molecules become truncated (Fig. 1B). Indeed, truncation may be a relatively rare event in the absence of NB-598, but the dramatically different stabilities of full-length and truncated SM lead to an equilibrium where their protein levels are comparable. Combined with the saturation of ERAD machinery, this may explain why overexpressed SM-V5 is less truncated than endogenous SM (Supplementary Fig. S3D) despite the two proteins having identical N-termini. Along similar lines, we previously found that overexpressed SM exhibits blunted cholesterol regulation [5].

**Figure 6.**
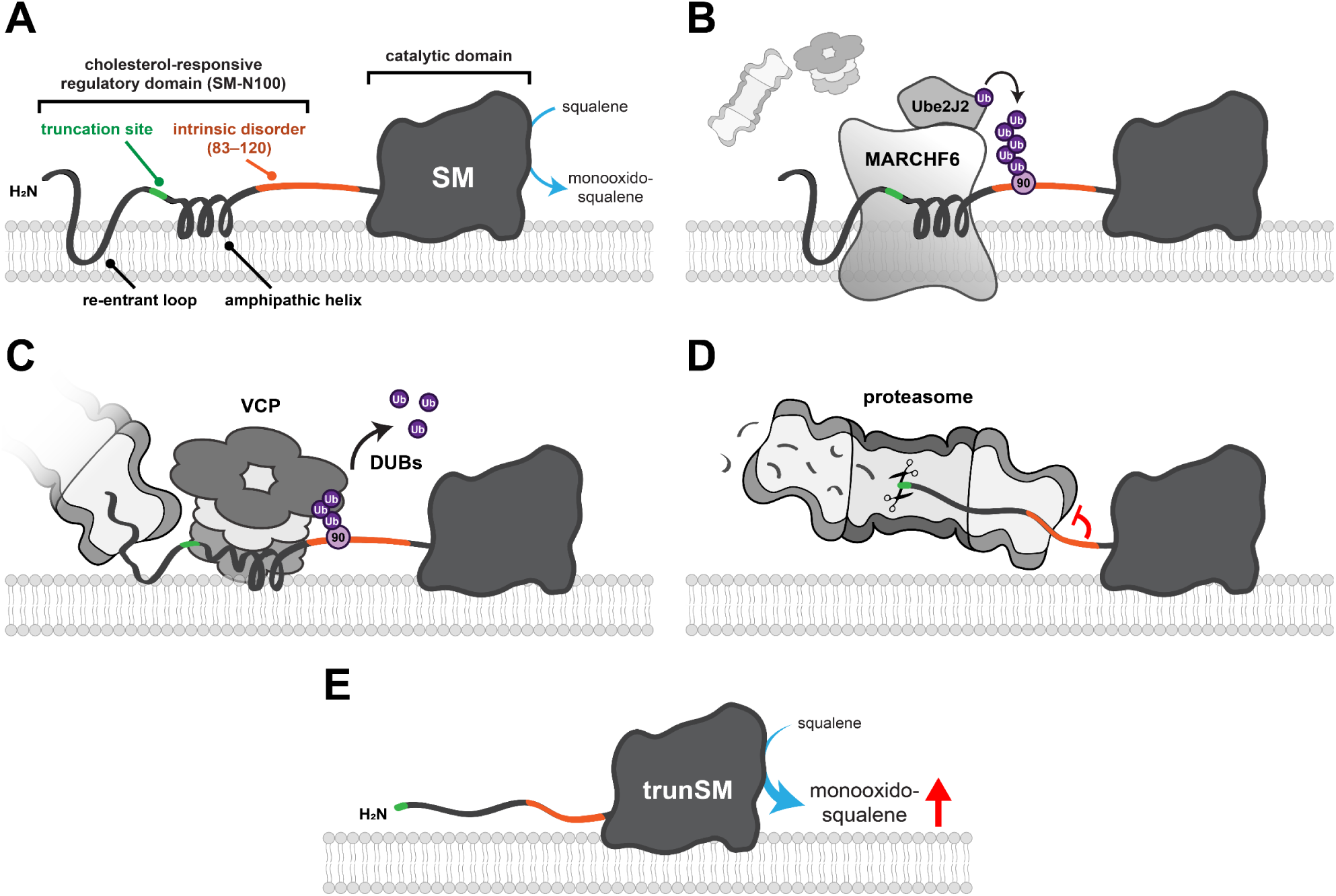
Model for the mechanism of SM truncation. **(A)** Full-length SM comprises the SM-N100 regulatory domain, containing a cholesterol-sensing re-entrant loop and amphipathic helix, and the C-terminal catalytic domain that converts squalene to monooxidosqualene. **(B)** Ube2J2 and MARCHF6 ubiquitinate the SM-N100 domain, likely at the known ubiquitination site Lys-90. **(C)** VCP is recruited to ubiquitinated SM and extracts the SM-N100 domain from the ER membrane, enabling the proteasome to begin degrading SM from its N-terminus. Deubiquitinases (DUBs) are required for this process. **(D)** The SM 81–120 disordered region impedes unfolding of the adjacent catalytic domain by the proteasome, preventing further degradation. **(E)** The undegraded portion of SM (trunSM) is released from the proteasome. The loss of part of the SM-N100 regulatory domain renders trunSM resistant to cholesterol-induced degradation, and therefore constitutively active.

The membrane association of trunSM (Fig. 5A) implies that the proteasome acts on SM directly at the ER without the need for cytosolic chaperones. While proteasomal recruitment to the ER has been described, this is generally in the context of interaction with the Sec61 translocon to export and degrade misfolded polypeptides [40]. One notable exception is the degradation of the yeast cadmium exporter Pca1p, in which interaction between the ubiquitinated substrate and the proteasome requires Doa10p (the orthologue of MARCHF6) and is bridged by Cdc48p (the orthologue of VCP) [41]. A similar pathway may be required for the truncation of SM, and presumably extends to its basal or even cholesterol-accelerated degradation. Precedent for the latter is found in the sterol-induced degradation of Hmg2p and its mammalian equivalent HMGCR (another rate-limiting enzyme of cholesterol synthesis), which also involves direct interaction with the proteasome at the ER membrane [41, 42]. In the case of SM, the AAA+ ATPase VCP likely provides the driving force to extract the membrane-associated components of the SM-N100 domain for degradation, although MARCHF6 may also contribute given the retrotranslocase function of Doa10p [43]. The role of deubiquitinases in SM truncation is less well-defined, but may involve the removal of ubiquitin chains from the Lys-82/90/100 cluster to facilitate processing by VCP or entry into the proteasome [44, 45]. Ubiquitination of two or more of these residues is seemingly required for maximal truncation (Fig. 3D), which may explain why a peptide containing Lys-90 alone was not enriched in a study that used VCP inhibition to accumulate and capture ERAD substrates [46]. Interestingly, proteasomal substrates with K63-linked ubiquitin chains are degraded less efficiently than those with the more typical K48-linkages [47], but whether SM is modified in this way is unknown.

### Structural determinants of truncation

In establishing SM as a substrate of partial proteasomal degradation, we identify the first eukaryotic enzyme known to be truncated in this manner. Of the few other reported substrates, almost all are soluble transcription factors and only two occur in mammalian cells: NFκB subunit p105 [24] and the Hedgehog signaling transducer Gli3 [25, 48]. Proteasomal truncation may be activating or inhibitory, depending on the substrate. In the case of NFκB, the proteasome degrades an inhibitory domain that sequesters the protein to the cytoplasm, thereby enabling its nuclear translocation [49]. Conversely, the transactivation domain of Gli3 is degraded to yield a dominant-negative repressor [25]. In yeast, partial degradation of Spt23p and Mga2p liberates them from the ER membrane to activate lipogenic gene expression, with Spt23p processing repressed by fatty acids [50]. Proteasomal processing can also have profound consequences at the organismal level: truncation of Gli3 or the *Drosophila* proteins Lola29M and Shavenbaby regulates processes including sex determination, differentiation, and stem cell maintenance [25, 28, 51, 52]. Like these examples, the truncation of SM eliminates a regulatory region (cholesterol-sensing elements of the SM-N100 domain) and yields a fragment with altered properties (constitutive activity and a peripheral membrane association).

For most truncation substrates, including NFκB [53], Gli3 [54] and Def1p [55], partial degradation requires a two-part signal: a low-complexity sequence (such as a glycine-or glutamine-rich region) and an adjacent tightly folded domain. The 19S regulatory particle of the proteasome is thought to poorly transduce an unfolding force when occupied by a low-complexity sequence, preventing disassembly of the folded domain and allowing substrate escape [27, 56]. However, we ruled out a low-complexity sequence as a major determinant of SM truncation (Supplementary Fig. S3A). Unique mechanisms have been described for the remaining substrates, with the exception of Lola29M where the structural features enabling truncation are unknown [51]. Spt23p and Mga2p are truncated upon proteasomal engagement at an internal hairpin loop, which leads to preferential degradation of the C-terminus rather than the tightly-folded N-terminal domain [29]. Such a mechanism is unlikely to control SM truncation, which is initiated at the N-terminus (Fig. 4D). In the case of Shavenbaby, a large (∼550 residue) intrinsically disordered domain comprising its degraded portion is juxtaposed with high predicted order in the adjacent, undegraded domain [28]. This most closely resembles SM, where truncation depends on the disordered 81–120 region (Fig. 4B), but the much shorter length of this region clearly represents a distinct signal.

On the other hand, the presence of a tightly folded and degradation-resistant domain is essential for all known examples of partial degradation. SM is no exception, and changes to the stability of its catalytic domain (*via* NB-598 binding or disruptive point mutations) are closely correlated with truncation (Fig. 1B and 4E). The split-domain structure of SM, in which substrate- and cofactor-binding regions are interspersed throughout the primary sequence [34], may contribute to a compact conformation that is unusually resistant to unfolding and degradation. Proximity of the catalytic domain to the ER membrane is presumably required for its interaction with the hydrophobic substrate squalene, and may sterically hinder engagement by VCP or the proteasome. Our findings thus reinforce the importance of a stable domain for disrupting proteasomal processivity, while also establishing a unique counterpart in the bipartite truncation signal: the 81–120 disordered region.

How does the 81–120 region promote truncation? The distance from the center of the proteasomal 20S core particle to the edge of the 19S regulatory particle is ∼200 Å [57], equivalent to ∼60 residues of a fully extended polypeptide chain. Therefore, as proteasomal degradation of SM reaches the predicted truncation site between residues 60 and 70 (Supplementary Fig. S2A), the 81–120 region is translocating through the regulatory particle and the catalytic domain is on the periphery of the complex. The conspicuous spacing of these three elements suggests that the 81–120 region is a proteasomal ‘stop signal’ that impedes further unfolding and incorporation of SM, analogous to the low-complexity sequences of other substrates (Fig. 6). This may explain why a minimum length of disorder is required for truncation (Fig. 4B), as well as the need for both the disordered region and catalytic domain stability. Without the former, the proteasome successfully transduces the force necessary to unfold the catalytic domain, and without the latter, proteasomal ATPases are still capable of disassembling the catalytic domain in their weakened state. In both cases, complete degradation is the result (Fig. 4B and 4E). It is noteworthy that the SM truncation site is unaltered when residues on the N-terminal side of the disorder are removed (Supplementary Fig. S3A) or when the disorder is duplicated (Fig. 4B), which supports the idea that the proteasome releases trunSM upon first encountering the 81–120 region.

Ubiquitination within residues 81–120 is unlikely to be solely responsible for its effect on truncation, given that the lowly truncated Δ91–110 mutant (Fig. 4B) retains both Lys-82 and Lys-90. Likewise, Lys-90 and Lys-100 are not conserved in highly truncated hamster and chicken orthologues (Fig. 4C). One possibility is that these sequences contain other lysine residues that serve as alternative ubiquitination sites. However, the number of lysine residues within orthologous regions (Supplementary Fig. S3B) does not correlate with the extent of truncation, and partial degradation is observed for a lamprey-derived sequence containing only one lysine residue. This strongly suggests that the intrinsically disordered nature of the region promotes truncation, but further work is required to elucidate the precise mechanism. Interestingly, the partial degradation of Spt23p and NF-κB is augmented by their homodimerization with a full-length counterpart, which protects the rescued fragment from complete proteolysis [58, 59]. Disordered regions often provide an interface for protein-protein interaction [60], and so a similar process may control SM truncation. It is currently unknown if SM forms dimers *in vivo*, but the transmembrane microprotein CASIMO1 is a confirmed interactor of SM [61] and GSK-3β was recently reported to associate with the isolated SM-N100 domain [62]. Given that GSK-3β is a soluble protein and much of SM-N100 is membrane-associated, the hydrophilic 81–100 region is a strong candidate for a binding site. Both CASIMO1 and GSK-3β are linked with metabolism [61, 63], warranting study into whether they impact on the production of the constitutively active trunSM.

### Consequences for cholesterol synthesis

Levels of full-length SM and trunSM are generally comparable across a range of human cell types (Fig. 1A), indicating that two enzyme pools are maintained: one which is subject to metabolic regulation and another which is constitutively active. As SM is a rate-limiting enzyme of cholesterol synthesis, this may establish a baseline level of pathway flux that can be fine-tuned depending on cholesterol availability. It is striking that amongst distant vertebrate orthologues of SM, there is strong structural conservation of the intrinsically disordered region (Supplementary Fig. S3B) and sequence conservation within the catalytic domain [5]. Truncation of SM occurs in CHO-7 cells (Supplementary Fig. S1) and may be characteristic of higher eukaryotes. Here, the basal activity of trunSM would permit a steady-state level of cholesterol synthesis to better meet the demands of multicellular organisms.

The presence of trunSM may also avert the complete ablation of SM activity under conditions of cholesterol excess, which would otherwise delay the eventual resumption of cholesterol synthesis or lead to dramatic accumulation of the substrate squalene [5]. Whilst previously considered a hydrocarbon intermediate with few biochemical properties, squalene is protective against cell death induced by lipid peroxidation [11] yet cytotoxic when it is unable to be effectively sequestered to lipid droplets [10, 64]. Increased squalene levels also accompany the dermal and gastrointestinal side effects of pharmacological SM inhibition in mammals [12]. Therefore, a persistent population of trunSM may be advantageous in clearing excess squalene and reducing its aberrant accumulation. Along similar lines, squalene itself stabilizes SM by binding to the SM-N100 domain and blunting its cholesterol-accelerated degradation [15]. It remains to be determined if squalene also influences the truncation of SM, given that both processes depend on MARCHF6 (Fig. 3C) [15]. It is also conceivable that trunSM is relevant in pathophysiological contexts, given that dysregulation of cholesterol synthesis is implicated in hepatocellular carcinoma [9] and prostate cancer [65]. However, this possibility awaits future investigation.

## Materials and methods

### Reagents and cell lines

Fetal calf serum (FCS), newborn calf serum (NCS), high-glucose Dulbecco’s Modified Eagle’s Medium (DMEM-HG), Roswell Park Memorial Institute 1640 medium (RPMI), DMEM/Ham’s Nutrient Mixture F-12 (DF-12), penicillin/streptomycin, Opti-MEM reduced serum medium, RNAiMAX transfection reagent, Lipofectamine 3000 transfection reagent, TRI reagent, and the SuperScript III First-Strand Synthesis kit were from Thermo-Fisher (Carlsbad, CA, US). Lipoprotein-deficient serum (LPDS) was prepared from NCS as described previously [8]. Primers, small interfering RNA (siRNA), protease inhibitor cocktail, and Tween-20 were from Sigma-Aldrich (St Louis, MO, US). The SensiMix SYBR No-ROX kit was from Bioline (London, UK). *SmaI* was from New England Biolabs (Ipswich, MA, US). Tris-glycine SDS-PAGE gels were prepared in-house. Immobilon Western chemiluminescent HRP substrate and nitrocellulose membranes were from Millipore (Burlington, MA, US). The QIAquick PCR purification kit was from Qiagen (Hilden, GE). *Trans*IT-2020 was from Mirus Bio (Madison, WA, US). Phosphate-buffered saline (PBS) was from UNSW (Sydney, AU). Skim milk powder was from Fonterra (Richmond, VIC, AU), and bovine serum albumin was from Bovogen Biologicals (Keilor, VIC, AU). Chemicals were from the following suppliers: cycloheximide (Sigma-Aldrich C7698), NB-598 (Chemscene CS-1274), cholesterol complexed with methyl-β-cyclodextrin (Chol/CD) (Sigma-Aldrich C4951), CB-5083 (Cayman Chemical Company 16276), MG132 (Sigma-Aldrich C2211), ALLN (Sigma-Aldrich A6185), PR-619 (Cayman Chemical Company 16276), WP1130 (Cayman Chemical Company 15277), ammonium chloride (Ajax Finechem 31), bafilomycin A1 (Sigma-Aldrich B1793), methotrexate (Cayman Chemical Company 13960), Tris-(hydroxy)methylamine (Ajax Finechem 2311), sodium chloride (Ajax Finechem 465); sodium dodecyl sulfate (Sigma-Aldrich 75746), magnesium chloride (Ajax Finechem 296), hydrochloric acid (Ajax Finechem 256), HEPES (Sigma-Aldrich 54457), potassium hydroxide (Ajax Finechem 405), potassium chloride (Ajax Finechem, 383), sodium ethylenediaminetetraacetic acid (Ajax Finechem 180), sodium ethylene glycol-bis(β-aminoethyl ether)-N,N,N′,N′-tetraacetic acid (Sigma-Aldrich E8145), sodium carbonate (Ajax Finechem 463), polyethyleneimine (Sigma-Aldrich 03880), glycine (Ajax Finechem 1083), methanol (Ajax Finechem 318), dimethyl sulfoxide (Ajax Finechem 2225), Ponceau S solution (Sigma-Aldrich P7170), glycerol (Ajax Finechem 242), bromophenol blue (Sigma-Aldrich B0126), β-mercaptoethanol (Sigma-Aldrich M3148).

HEK293T cells were a gift from the UNSW School of Medical Sciences (UNSW, Sydney NSW, Australia), HepG2 and Huh7 cells were gifts from the Centre for Cardiovascular Research (UNSW, Sydney NSW, Australia), Be(2)-C and HeLaT cells were gifts from Drs. Louise Lutze-Mann and Noel Whitaker (UNSW, Sydney NSW, Australia), and CHO-7 cells were a gift from Drs. Joseph Goldstein and Michael Brown (UT Southwestern Medical Center, Dallas TX, USA).

### Cell culture

Cells were maintained in a humidified incubator at 37°C and 5% CO_2_ in maintenance medium (DMEM-HG [HEK293T, HepG2, Huh7, Be(2)-C], RPMI [HeLaT] or DF-12 [CHO-7]; 10% [v/v] FCS for human cells or 5% [v/v] LPDS [50 mg/mL protein] for CHO-7 cells; 100 U/mL penicillin; 100 μg/mL streptomycin). To improve HEK293T and HepG2 surface adhesion, culture vessels were treated with 25 μg/mL polyethyleneimine in phosphate-buffered saline (PBS) for 15 min prior to cell seeding. Plasmid and siRNA transfections were performed in maintenance medium lacking penicillin and streptomycin. For all treatments, appropriate solvent controls were used (water [Chol/CD, ammonium chloride]; dimethyl sulfoxide [cycloheximide, NB-598, CB-5083, MG132, ALLN, bafilomycin A1, PR-619, WP1130, methotrexate]) and the final concentration of dimethyl sulfoxide did not exceed 0.2% (v/v) in cell culture medium. Treatments were delivered in full medium refreshes, and all experiments were a total of 72 h in duration.

### Plasmids

A pcDNA3.1/V5-His TOPO expression vector (Invitrogen) encoding the protein-coding sequence of human SM (NM_003129.4) fused with three N-terminal HA tags, a C-terminal linker sequence, and C-terminal V5 and 6 × His tags ([HA]_3_-SM-V5) was generated previously in our laboratory by Dr Julian Stevenson. Codon-optimized nucleotide sequences encoding orthologues of human SM-N100 were previously obtained from GenScript [7], and sequences encoding orthologues of human SM residues 101–120 were derived using the Integrated DNA Technologies codon optimization tool. Domain insertions and deletions within the (HA)_3_-SM-V5 construct were generated using the polymerase-incomplete primer extension cloning method and sequence- and ligation-independent cloning method [5, 6], and domain and nucleotide substitutions were generated using the overlap extension cloning method [4], as described previously [9]. To generate standards for the absolute quantification of mRNA levels, qRT-PCR products were amplified from HEK293T cDNA and inserted into the pGL3-Basic vector (Promega) using the overlap extension cloning method [4]. The identity of all plasmids was confirmed *via* Sanger dideoxy sequencing. The plasmids used in this study are listed in Table S1, and the primer sequences used for DNA cloning are listed in Table S2.

### siRNA and plasmid transfection

To downregulate gene expression or transiently overexpress SM-derived constructs, cells were seeded into 12-well plates. The next day, cells were transfected with 15 pmol siRNA using RNAiMAX (Invitrogen; 15 pmol siRNA : 2 µL reagent) or 1 µg expression vector using Lipofectamine 3000 (Invitrogen; 1 µg DNA : 2 µL reagent with 2 µL P3000 supplemental reagent), delivered in Opti-MEM. After 24 h, cells were refreshed in maintenance medium, treated as specified in figure legends, and harvested 48 h after transfection. The siRNAs used in this study are listed in Table S1.

### Protein harvest, SDS-PAGE and immunoblotting

To quantify protein levels, cells were seeded into 12-well plates and treated as specified in figure legends. Total protein was harvested in 2% SDS lysis buffer (10 mM Tris-HCl [pH 7.6], 2% [w/v] SDS, 100 mM NaCl) containing 2% [v/v] protease inhibitor cocktail, passed 20 times through a 21-gauge needle, and vortexed for 20 min. Lysate protein content was quantified using the bicinchoninic acid assay (Thermo-Fisher), and sample concentrations were normalized by dilution in 2% SDS lysis buffer and 1× Laemmli buffer (50 mM Tris-HCl [pH 6.8], 2% [w/v] SDS, 5% [v/v] glycerol, 0.04% [w/v] bromophenol blue, 1% [v/v] β-mercaptoethanol). Samples were heated at 95°C for 5 min and separated by 10% (w/v) Tris-glycine SDS-PAGE, unless otherwise specified in figure legends. Proteins were electroblotted onto nitrocellulose membranes and blocked in 5% skim milk powder (Diploma) in PBS with 0.1% (v/v) Tween-20 (PBST), or in 5% bovine serum albumin in PBST for FLAG detection. Immunoblotting was performed using rabbit polyclonal anti-SM(SQLE) (Proteintech 12544-1-AP; 1:2,500 at 4°C for 16 h), rabbit monoclonal anti-GAPDH (Cell Signaling Technology 2118; 1:2,500 at 4°C for 16 h), rabbit monoclonal anti-HA (Cell Signaling Technology 3724; 1:2,000 at 4°C for 16 h), mouse monoclonal anti-V5 (Invitrogen R960-25; 1:5,000 at room temperature for 1 h), rabbit polyclonal anti-FLAG (Millipore F7425; 1:10,000 at 4°C for 16 h), IRDye 680RD donkey anti-rabbit IgG (LI-COR Biosciences LCR-926-68073; 1:5,000 [SM detection] or 1:10,000 at room temperature for 1 h), IRDye 800CW donkey anti-mouse IgG (LI-COR Biosciences LCR-926-32212; 1:10,000 at room temperature for 1 h), peroxidase-conjugated AffiniPure donkey anti-rabbit IgG (Jackson ImmunoResearch Laboratories 711-035-152; 1:10,000 at room temperature for 1 h), and peroxidase-conjugated AffiniPure donkey anti-mouse IgG (Jackson ImmunoResearch Laboratories 715-035-150; 1:10,000 at room temperature for 1 h). All antibodies were diluted in 5% bovine serum albumin in PBST, except for anti-FLAG and peroxidase-conjugated antibodies, which were diluted in 5% skim milk in PBST. Fluorescence-based detection of SM, GAPDH, HA and V5 was performed using an Odyssey Clx imager (LI-COR Biosciences), and enhanced chemiluminescence-based detection of FLAG was performed using Immobilon Western chemiluminescent HRP substrate (Millipore) and an ImageQuant LAS 500 imager (Cytiva Life Sciences). Due to low protein levels following differential solubilization of microsomal membranes, enhanced chemiluminescence was used to detect FLAG and V5 in these samples. Densitometry analysis of fluorescence images was performed using Image Studio Lite v5.2.5 (LI-COR Biosciences).

### RNA harvest and qRT-PCR

To quantify *squalene epoxidase* (*SQLE*) gene expression, cells were seeded in triplicate into 12-well plates and transfected with siRNA as specified in figure legends. Total RNA was harvested using TRI reagent (Sigma-Aldrich) and polyadenylated RNA was reverse transcribed using the SuperScript III First Strand Synthesis kit (Invitrogen). cDNA products were used as the template for quantitative reverse transcription-PCR (qRT-PCR) using the SensiMix SYBR No-ROX kit (Bioline). For relative quantification of gene expression, mRNA levels were normalized to the *porphobilinogen deaminase* (*PBGD*) housekeeping gene using the comparative C_T_ method [10] and adjusted relative to the control siRNA condition, as specified in figure legends. For absolute quantification of *SQLE* expression, plasmids containing qPCR amplicon sequences were linearized by digestion with *SmaI* for 1 h and purified using the QIAquick PCR purification kit (Qiagen). Linearized plasmids were quantified using spectrophotometry, serially diluted in nuclease-free water to concentrations of between ∼5×10^2^ and ∼5×10^8^ target copies/µL and used as the template for qPCR in triplicate as described above. A standard curve of log(target sequence copies) vs. C_T_ value was generated and compared with C_T_ values from cDNA samples to quantify gene expression. Data were expressed in units of cDNA molecules per µg of reverse transcribed RNA. The primer sequences used for qRT-PCR in this study are listed in Table S2.

### Cell fractionation and differential solubilization

To examine protein membrane association, cells were seeded into 10 cm dishes and treated as specified in figure legends. Microsomal membranes were isolated as described in [11] with some modifications. Briefly, cells were scraped in cold PBS, pelleted at 1,000 × *g* and 4°C for 5 min, and lysed in 500 µl buffer F1 (10 mM HEPES-KOH [pH 7.4], 10 mM KCl, 1.5 mM MgCl_2_, 5 mM sodium EDTA, 5 mM sodium EGTA, 250 mM sucrose, 2% [v/v] protease inhibitor cocktail). Lysates were centrifuged at 1,000 × *g* and 4°C for 10 min, and the supernatant was centrifuged at 20,000 × *g* and 4°C for 30 min. The 20,000 × *g* supernatant was collected and designated the cytosolic fraction. The 20,000 × *g* pellet was resuspended in 100 µl buffer F2 (10 mM Tris-HCl [pH 7.4], 100 mM NaCl, 1% [w/v] SDS, 2% [v/v] protease inhibitor cocktail) and designated the membrane fraction. Protein content was quantified using the bicinchoninic acid assay (Thermo-Fisher), and sample concentrations were normalized by dilution in buffer F1 or buffer F2, plus 1× Laemmli buffer, for immunoblotting analysis.

To determine the peripheral or integral nature of protein membrane association, differential solubilization of microsomal membranes was performed as described in [11] with some modifications. Briefly, cells were seeded into 14.5 cm dishes and transfected with 40 µg pTK-SM-N100-GFP-V5 expression vector using *TransIT*-2020 (Mirus Bio; 1 µg DNA: 2 µL reagent), delivered in Opti-MEM. After 24 h, cells were refreshed in maintenance medium for a further 24 h, and microsomal membranes were isolated as described above. Equivalent volumes of membrane preparations (20 µl) were treated with 200 µl buffer F1, 1% (w/v) SDS (with 10 mM Tris-HCl [pH 7.4]), 0.1 M Na_2_CO_3_ (pH 11.5), or 1 M NaCl (with 10 mM Tris-HCl [pH 7.4]), and incubated at 4°C with end-over-end mixing for 30 min. Mixtures were then centrifuged at 20,000 × *g* and 4°C for 30 min. The soluble supernatant fraction was collected, and the insoluble pellet fraction was resuspended in 200 µl buffer F3 (buffer F1 containing 100 mM NaCl). Equal volumes of supernatant and pellet fractions were mixed with 1× Laemmli buffer for immunoblotting analysis.

### Sequences and alignments

DNA sequences of protein-coding *SQLE* isoforms (*fullSQLE*, NM_003129.4; *trunSQLE1*, ENST00000523430.5; *trunSQLE2*, XM_011517246.2) were obtained from the RefSeq-(GRCh38.p13 109.20200228) and GENCODE-annotated (GRCh38.p13 GCA_000001405.28) human genomes. Protein sequences of human SM (*Homo sapiens*, Q14534), Chinese hamster SM (*Cricetulus griseus*, A0A3L7IPT3), chicken SM (*Gallus gallus*, A0A1D5NWK3), zebrafish SM (*Danio rerio*, F1QDN5), sea lamprey SM (*Petromyzon marinus*, S4R6S3) and yeast Erg1p (*Saccharomyces cerevisiae*, P32476) were obtained from the UniProt database [12]. Protein sequence complexity was predicted using the SEG [13], CAST [14] and fLPS [15] algorithms, and regions identified by all three tools were defined as low-complexity sequences. Protein intrinsic disorder was predicted using the online tools SPOT-dis2 [16], MFDp2 [17], AUCpreD [18], IUPred2A [19], DISOPRED3 [20], PrDOS [21] and DisProt (VL2E) [22], and residues with an average intrinsic disorder probability of >0.5 were defined as intrinsically disordered. Protein sequence alignments were generated using Geneious Basic v2020.1 (Biomatters Ltd.) with a BLOSUM62 cost matrix.

### Data analysis and presentation

Data were normalized as described in figure legends. Data visualization and statistical testing were performed using GraphPad Prism v8.4 (GraphPad Software Inc.) as specified in figure legends. Thresholds for statistical significance were defined as: *, *p* ≤ 0.05; **, *p* ≤ 0.01. Schematics and figures were assembled using Adobe Illustrator v24.1 (Adobe Inc.).

## Acknowledgements

We thank Dr Julian Stevenson for generating the pCMV-(HA)_3_-SM-V5 plasmid used in this study, Dr Ngee Kiat (Jake) Chua for insightful discussions, and the members of the Brown laboratory for critically reviewing this manuscript. This work was supported by Australian Research Council Grant DP170101178 and a NSW Health Investigator Development Grant. H.W.C. is a recipient of an Australian Research Training Program scholarship.

## Competing interests

The authors declare that there are no competing interests associated with the manuscript.

## Supplementary data

### Supplementary figures

**Figure S1. Related to Fig. 1.**
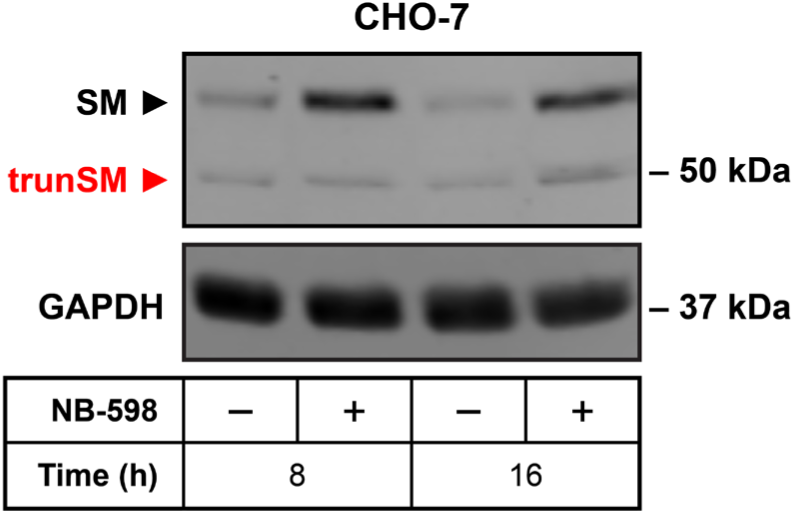
CHO-7 cells express a trunSM-like protein. CHO-7 cells were treated in the presence or absence of 1 µM NB-598 for the indicated times, and immunoblotting was performed for SM and truncated SM (trunSM, red). Immunoblot is representative of *n* = 2 independent experiments.

**Figure S2. Related to Fig. 3.**
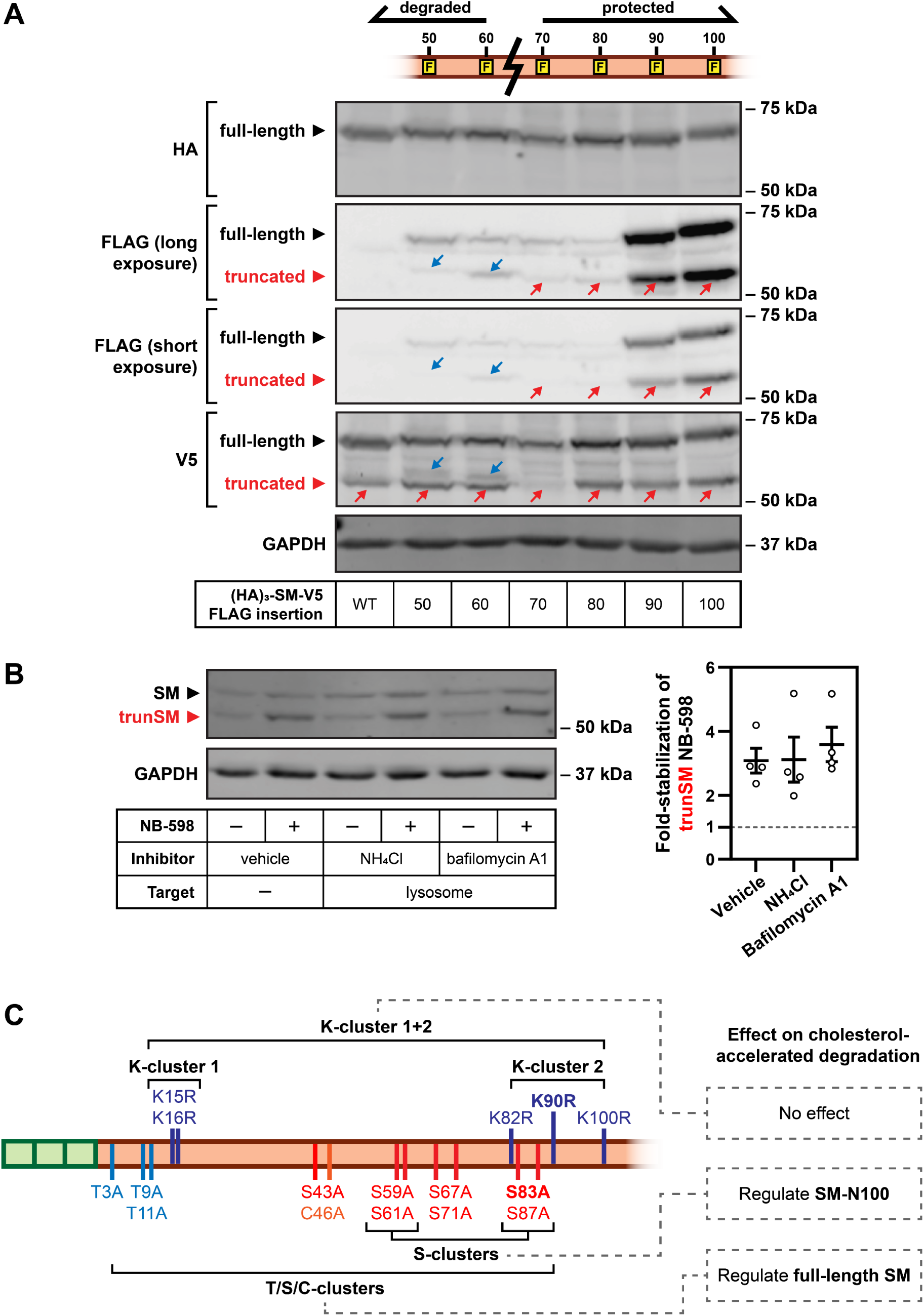
**(A)** The estimated SM truncation site is between residues 60 and 70. HEK293T cells were transfected with the indicated constructs for 24 h, treated with 1 µM NB-598 for 16 h, and then treated with 20 µM MG132 for 8 h. Immunoblot is representative of *n* ≥ 2 independent experiments. Red arrows indicate fragments corresponding to trunSM, and blue arrows indicate additional FLAG-tagged fragments that do not correspond to trunSM. **(B)** SM truncation does not depend on the lysosome. HEK293T cells were treated with 20 mM ammonium chloride (NH_4_Cl) or 10 nM bafilomycin A1, in the presence or absence of 1 µM NB-598, for 8 h. Graph depicts densitometric quantification of trunSM stabilization by NB-598. Data presented as mean ± SEM from *n* = 4 independent experiments. **(C)** Schematic of putative ubiquitination sites within the SM-N100 domain. Lysine residues are not required for cholesterol-accelerated degradation of SM or SM-N100 [1, 2]. Serine residues are required for maximal cholesterol-accelerated degradation of SM-N100 [1], while clusters of threonine, cysteine and serine residues are required for maximal cholesterol-accelerated degradation of full-length SM [1]. Bolded residues indicate known ubiquitination sites [1, 3].

**Figure S3. Related to Fig. 4.**
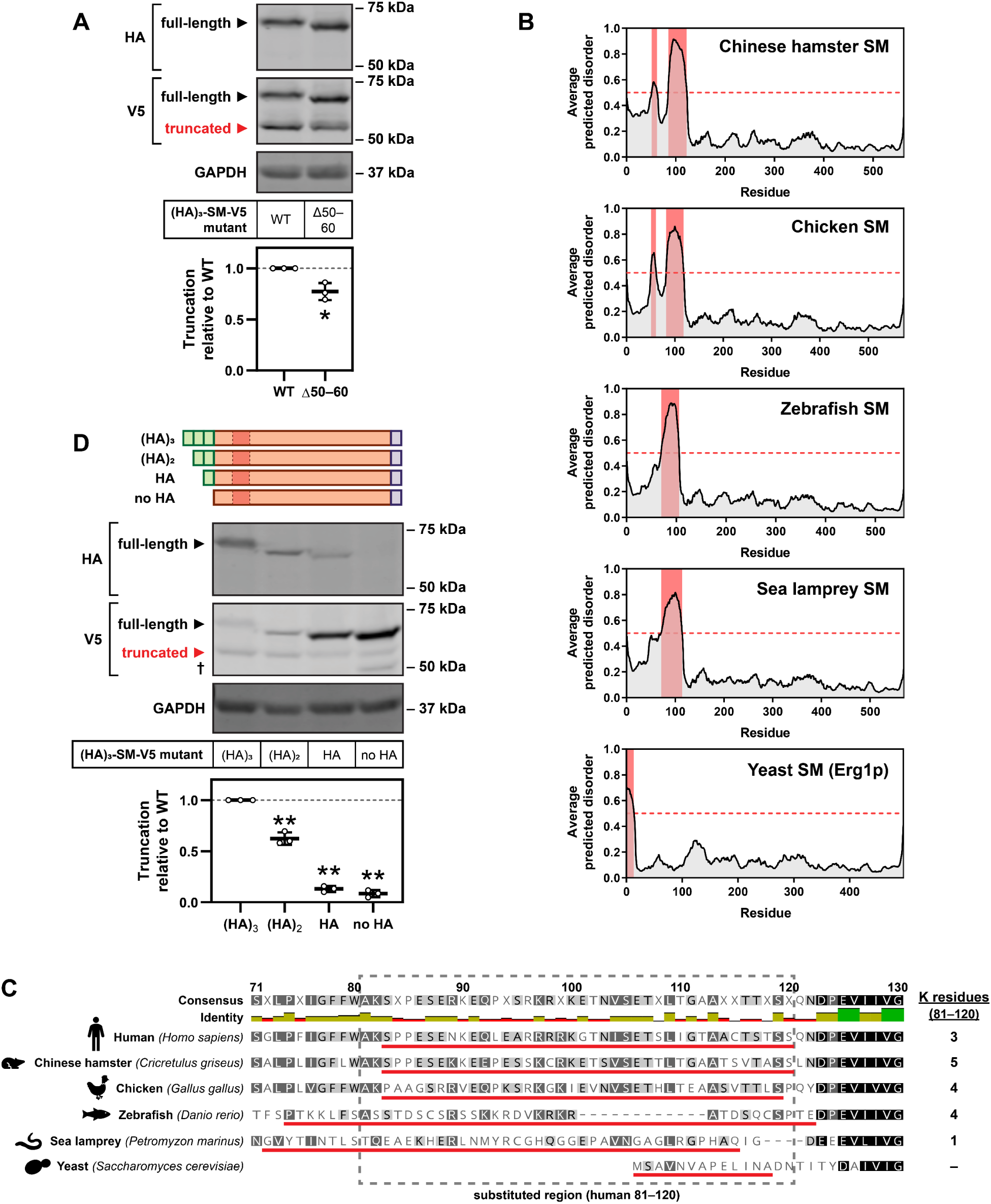
**(A)** A low-complexity sequence within the SM-N100 domain has a small effect on SM truncation. HEK293T cells were transfected with the indicated constructs for 24 h and refreshed in maintenance medium for 24 h. Graph depicts densitometric quantification of truncation normalized to the wild-type (WT) construct, which was set to 1 (dotted line). Data presented as mean ± SEM from *n* = 3 independent experiments (*, p ≤ 0.05, two-tailed paired t-test vs. WT). **(B)** The intrinsic disorder of the SM 81–120 region is highly conserved amongst SM orthologues. Intrinsically disordered regions (red) are indicated for SM orthologues from Chinese hamster (*Cricetulus griseus*), chicken (*Gallus gallus*), zebrafish (*Danio rerio*), sea lamprey (*Petromyzon marinus*) and yeast (*Saccharomyces cerevisiae*). **(C)** The sequence of the SM 81–120 region is poorly conserved amongst SM orthologues. Alignment of human SM residues 71–130 with SM orthologues from the indicated species. Red bars indicate regions of intrinsic disorder, and grey dashed box indicates regions that were substituted into SM constructs in Fig. 4C. **(D)** Removal of HA tags from the SM N-terminus reduces truncation. HEK293T cells were transfected with the indicated constructs for 24 h and refreshed in maintenance medium for 24 h. Graph depicts densitometric quantification of truncation normalized to the wild-type (WT) construct, which was set to 1 (dotted line). Data presented as mean ± SEM from *n* = 3 independent experiments (**, p ≤ 0.01, two-tailed paired t-test vs. WT). Dagger indicates a non-trunSM fragment.

### Supplementary tables

**Table S1.**
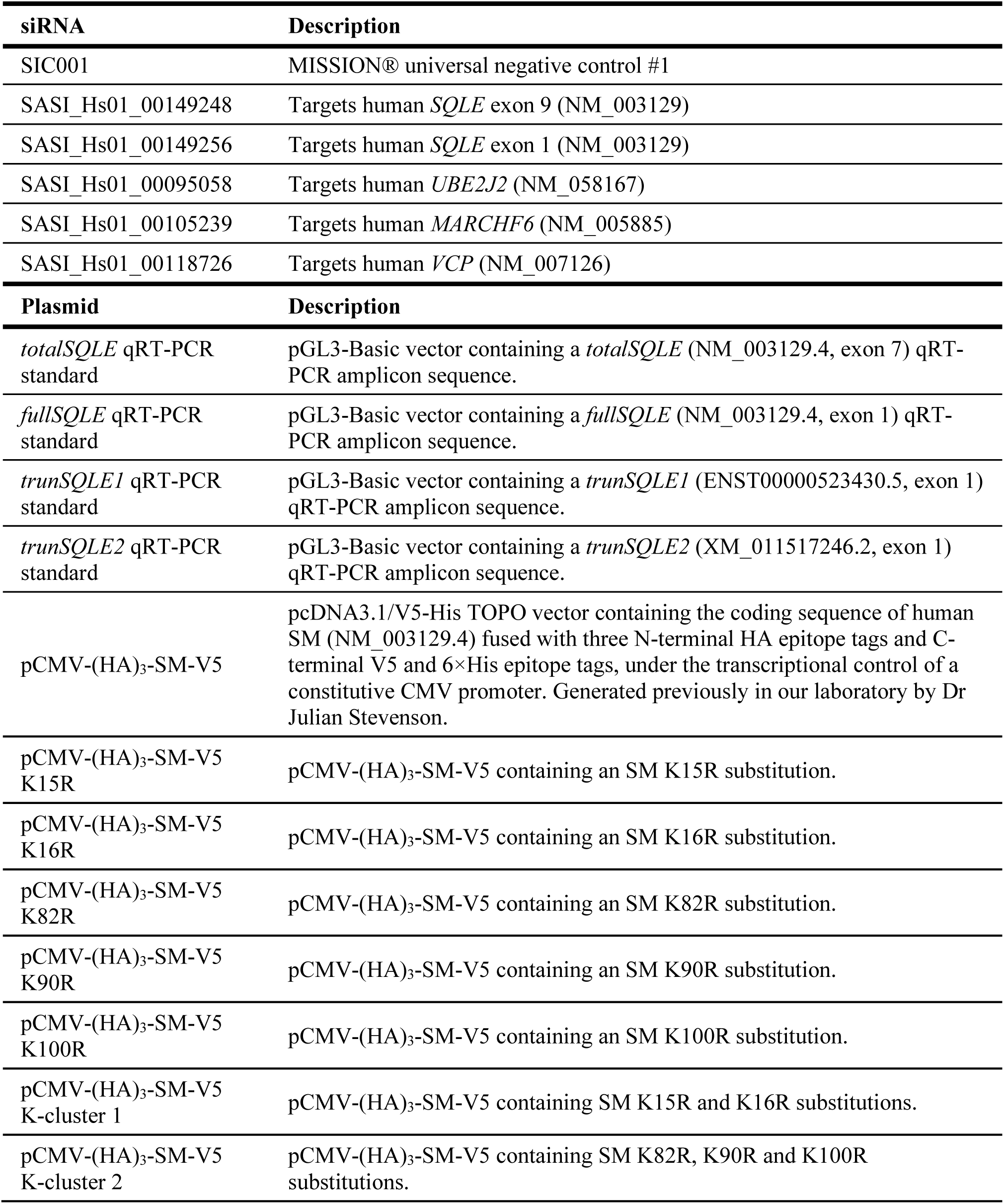
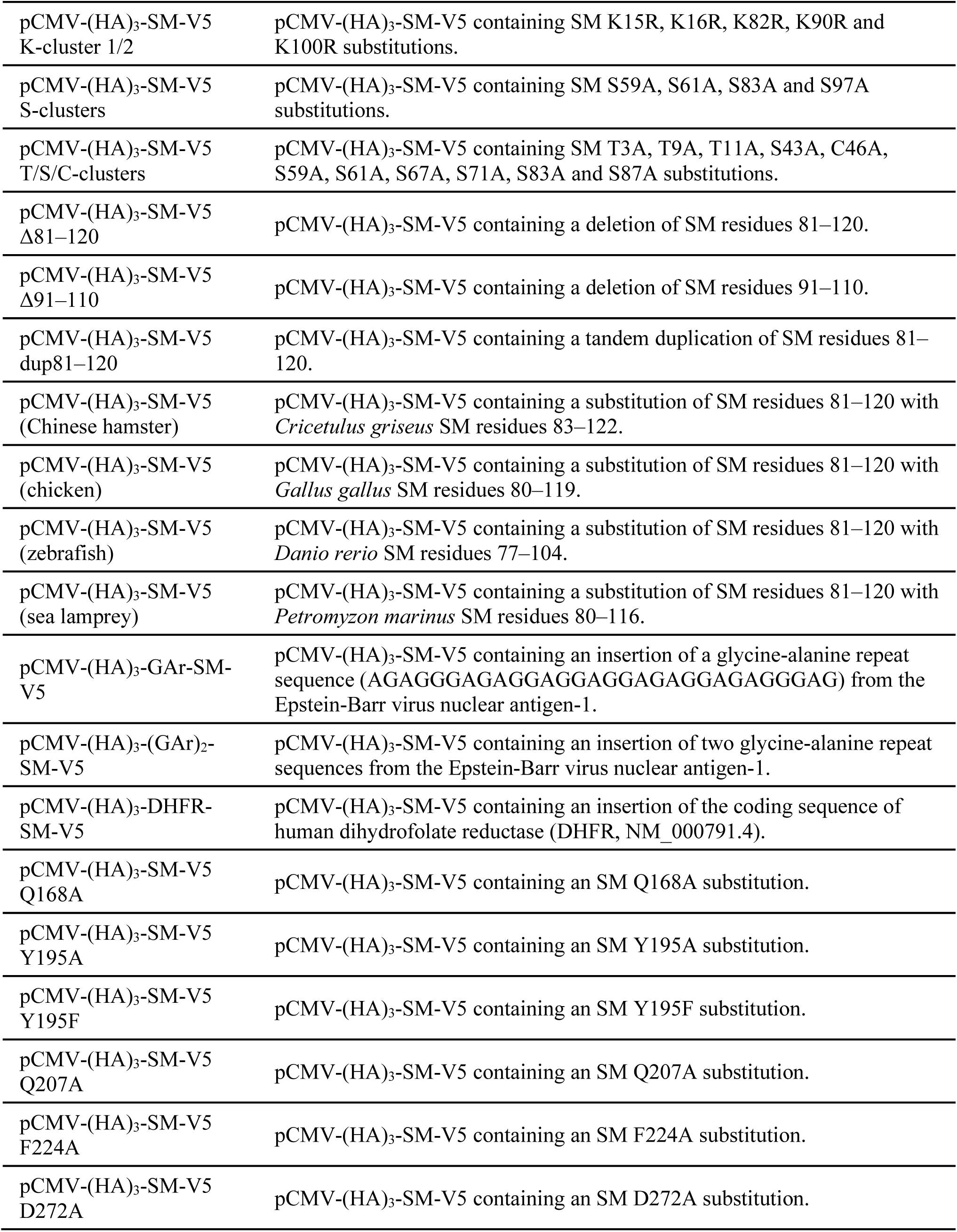
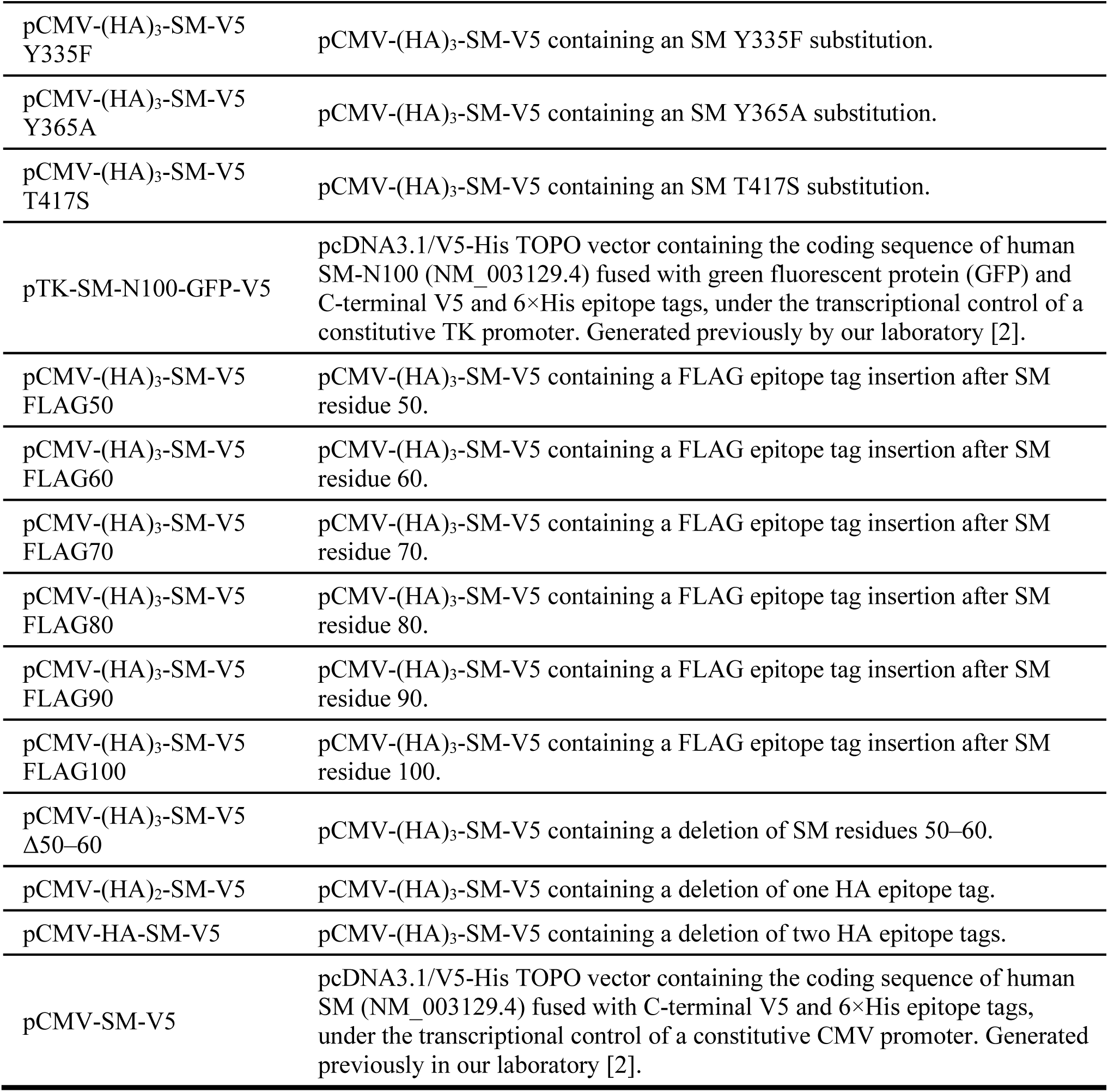
siRNA and plasmids used for transfection.

| siRNA | Description |
| --- | --- |
| SIC001 | MISSION® universal negative control #1 |
| SASI_Hs01_00149248 | Targets human *SQLE* exon 9 (NM_003129) |
| SASI_Hs01_00149256 | Targets human *SQLE* exon 1 (NM_003129) |
| SASI_Hs01_00095058 | Targets human *UBE2J2* (NM_058167) |
| SASI_Hs01_00105239 | Targets human *MARCHF6* (NM_005885) |
| SASI_Hs01_00118726 | Targets human *VCP* (NM_007126) |
| Plasmid | Description |
| *totalSQLE* qRT-PCR standard | pGL3-Basic vector containing a *totalSQLE* (NM_003129.4, exon 7) qRT-PCR amplicon sequence. |
| *fullSQLE* qRT-PCR standard | pGL3-Basic vector containing a *fullSQLE* (NM_003129.4, exon 1) qRT-PCR amplicon sequence. |
| *trunSQLE1* qRT-PCR standard | pGL3-Basic vector containing a *trunSQLE1* (ENST00000523430.5, exon 1) qRT-PCR amplicon sequence. |
| *trunSQLE2* qRT-PCR standard | pGL3-Basic vector containing a *trunSQLE2* (XM_011517246.2, exon 1) qRT-PCR amplicon sequence. |
| pCMV-(HA)<sub>3</sub>-SM-V5 | pcDNA3.1/V5-His TOPO vector containing the coding sequence of human SM (NM_003129.4) fused with three N-terminal HA epitope tags and C-terminal V5 and 6×His epitope tags, under the transcriptional control of a constitutive CMV promoter. Generated previously in our laboratory by Dr Julian Stevenson. |
| pCMV-(HA)<sub>3</sub>-SM-V5 K15R | pCMV-(HA)<sub>3</sub>-SM-V5 containing an SM K15R substitution. |
| pCMV-(HA)<sub>3</sub>-SM-V5 K16R | pCMV-(HA)<sub>3</sub>-SM-V5 containing an SM K16R substitution. |
| pCMV-(HA)<sub>3</sub>-SM-V5 K82R | pCMV-(HA)<sub>3</sub>-SM-V5 containing an SM K82R substitution. |
| pCMV-(HA)<sub>3</sub>-SM-V5 K90R | pCMV-(HA)<sub>3</sub>-SM-V5 containing an SM K90R substitution. |
| pCMV-(HA)<sub>3</sub>-SM-V5 K100R | pCMV-(HA)<sub>3</sub>-SM-V5 containing an SM K100R substitution. |
| pCMV-(HA)<sub>3</sub>-SM-V5 K-cluster 1 | pCMV-(HA)<sub>3</sub>-SM-V5 containing SM K15R and K16R substitutions. |
| pCMV-(HA)<sub>3</sub>-SM-V5 K-cluster 2 | pCMV-(HA)<sub>3</sub>-SM-V5 containing SM K82R, K90R and K100R substitutions. |
| pCMV-(HA)<sub>3</sub>-SM-V5<br>K-cluster 1/2 | pCMV-(HA)<sub>3</sub>-SM-V5 containing SM K15R, K16R, K82R, K90R and K100R substitutions. |
| pCMV-(HA)<sub>3</sub>-SM-V5<br>S-clusters | pCMV-(HA)<sub>3</sub>-SM-V5 containing SM S59A, S61A, S83A and S97A substitutions. |
| pCMV-(HA)<sub>3</sub>-SM-V5<br>T/S/C-clusters | pCMV-(HA)<sub>3</sub>-SM-V5 containing SM T3A, T9A, T11A, S43A, C46A, S59A, S61A, S67A, S71A, S83A and S87A substitutions. |
| pCMV-(HA)<sub>3</sub>-SM-V5<br>Δ81–120 | pCMV-(HA)<sub>3</sub>-SM-V5 containing a deletion of SM residues 81–120. |
| pCMV-(HA)<sub>3</sub>-SM-V5<br>Δ91–110 | pCMV-(HA)<sub>3</sub>-SM-V5 containing a deletion of SM residues 91–110. |
| pCMV-(HA)<sub>3</sub>-SM-V5<br>dup81–120 | pCMV-(HA)<sub>3</sub>-SM-V5 containing a tandem duplication of SM residues 81–120. |
| pCMV-(HA)<sub>3</sub>-SM-V5<br>(Chinese hamster) | pCMV-(HA)<sub>3</sub>-SM-V5 containing a substitution of SM residues 81–120 with *Cricetulus griseus* SM residues 83–122. |
| pCMV-(HA)<sub>3</sub>-SM-V5<br>(chicken) | pCMV-(HA)<sub>3</sub>-SM-V5 containing a substitution of SM residues 81–120 with *Gallus gallus* SM residues 80–119. |
| pCMV-(HA)<sub>3</sub>-SM-V5<br>(zebrafish) | pCMV-(HA)<sub>3</sub>-SM-V5 containing a substitution of SM residues 81–120 with *Danio rerio* SM residues 77–104. |
| pCMV-(HA)<sub>3</sub>-SM-V5<br>(sea lamprey) | pCMV-(HA)<sub>3</sub>-SM-V5 containing a substitution of SM residues 81–120 with *Petromyzon marinus* SM residues 80–116. |
| pCMV-(HA)<sub>3</sub>-GAR-SM-V5 | pCMV-(HA)<sub>3</sub>-SM-V5 containing an insertion of a glycine-alanine repeat sequence (AGAGGGAGAGGAGGAGGAGAGGGAG) from the Epstein-Barr virus nuclear antigen-1. |
| pCMV-(HA)<sub>3</sub>-(GAR)<sub>2</sub>-SM-V5 | pCMV-(HA)<sub>3</sub>-SM-V5 containing an insertion of two glycine-alanine repeat sequences from the Epstein-Barr virus nuclear antigen-1. |
| pCMV-(HA)<sub>3</sub>-DHFR-SM-V5 | pCMV-(HA)<sub>3</sub>-SM-V5 containing an insertion of the coding sequence of human dihydrofolate reductase (DHFR, NM_000791.4). |
| pCMV-(HA)<sub>3</sub>-SM-V5<br>Q168A | pCMV-(HA)<sub>3</sub>-SM-V5 containing an SM Q168A substitution. |
| pCMV-(HA)<sub>3</sub>-SM-V5<br>Y195A | pCMV-(HA)<sub>3</sub>-SM-V5 containing an SM Y195A substitution. |
| pCMV-(HA)<sub>3</sub>-SM-V5<br>Y195F | pCMV-(HA)<sub>3</sub>-SM-V5 containing an SM Y195F substitution. |
| pCMV-(HA)<sub>3</sub>-SM-V5<br>Q207A | pCMV-(HA)<sub>3</sub>-SM-V5 containing an SM Q207A substitution. |
| pCMV-(HA)<sub>3</sub>-SM-V5<br>F224A | pCMV-(HA)<sub>3</sub>-SM-V5 containing an SM F224A substitution. |
| pCMV-(HA)<sub>3</sub>-SM-V5<br>D272A | pCMV-(HA)<sub>3</sub>-SM-V5 containing an SM D272A substitution. |
| pCMV-(HA)<sub>3</sub>-SM-V5<br>Y335F | pCMV-(HA)<sub>3</sub>-SM-V5 containing an SM Y335F substitution. |
| pCMV-(HA)<sub>3</sub>-SM-V5<br>Y365A | pCMV-(HA)<sub>3</sub>-SM-V5 containing an SM Y365A substitution. |
| pCMV-(HA)<sub>3</sub>-SM-V5<br>T417S | pCMV-(HA)<sub>3</sub>-SM-V5 containing an SM T417S substitution. |
| pTK-SM-N100-GFP-V5 | pcDNA3.1/V5-His TOPO vector containing the coding sequence of human SM-N100 (NM_003129.4) fused with green fluorescent protein (GFP) and C-terminal V5 and 6×His epitope tags, under the transcriptional control of a constitutive TK promoter. Generated previously by our laboratory [2]. |
| pCMV-(HA)<sub>3</sub>-SM-V5<br>FLAG50 | pCMV-(HA)<sub>3</sub>-SM-V5 containing a FLAG epitope tag insertion after SM residue 50. |
| pCMV-(HA)<sub>3</sub>-SM-V5<br>FLAG60 | pCMV-(HA)<sub>3</sub>-SM-V5 containing a FLAG epitope tag insertion after SM residue 60. |
| pCMV-(HA)<sub>3</sub>-SM-V5<br>FLAG70 | pCMV-(HA)<sub>3</sub>-SM-V5 containing a FLAG epitope tag insertion after SM residue 70. |
| pCMV-(HA)<sub>3</sub>-SM-V5<br>FLAG80 | pCMV-(HA)<sub>3</sub>-SM-V5 containing a FLAG epitope tag insertion after SM residue 80. |
| pCMV-(HA)<sub>3</sub>-SM-V5<br>FLAG90 | pCMV-(HA)<sub>3</sub>-SM-V5 containing a FLAG epitope tag insertion after SM residue 90. |
| pCMV-(HA)<sub>3</sub>-SM-V5<br>FLAG100 | pCMV-(HA)<sub>3</sub>-SM-V5 containing a FLAG epitope tag insertion after SM residue 100. |
| pCMV-(HA)<sub>3</sub>-SM-V5<br>Δ50–60 | pCMV-(HA)<sub>3</sub>-SM-V5 containing a deletion of SM residues 50–60. |
| pCMV-(HA)<sub>2</sub>-SM-V5 | pCMV-(HA)<sub>3</sub>-SM-V5 containing a deletion of one HA epitope tag. |
| pCMV-HA-SM-V5 | pCMV-(HA)<sub>3</sub>-SM-V5 containing a deletion of two HA epitope tags. |
| pCMV-SM-V5 | pcDNA3.1/V5-His TOPO vector containing the coding sequence of human SM (NM_003129.4) fused with C-terminal V5 and 6×His epitope tags, under the transcriptional control of a constitutive CMV promoter. Generated previously in our laboratory [2]. |

**Table S2.**
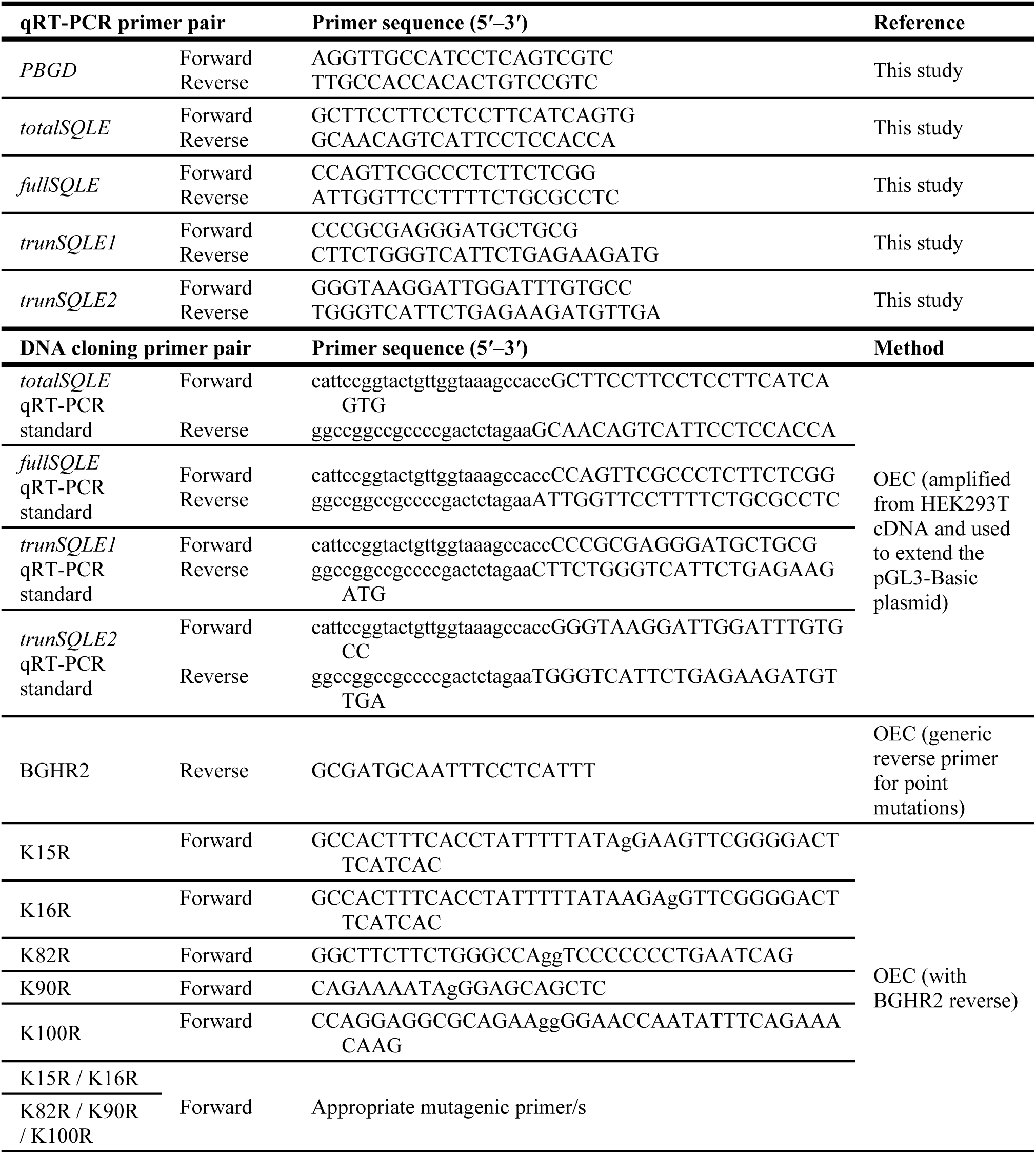
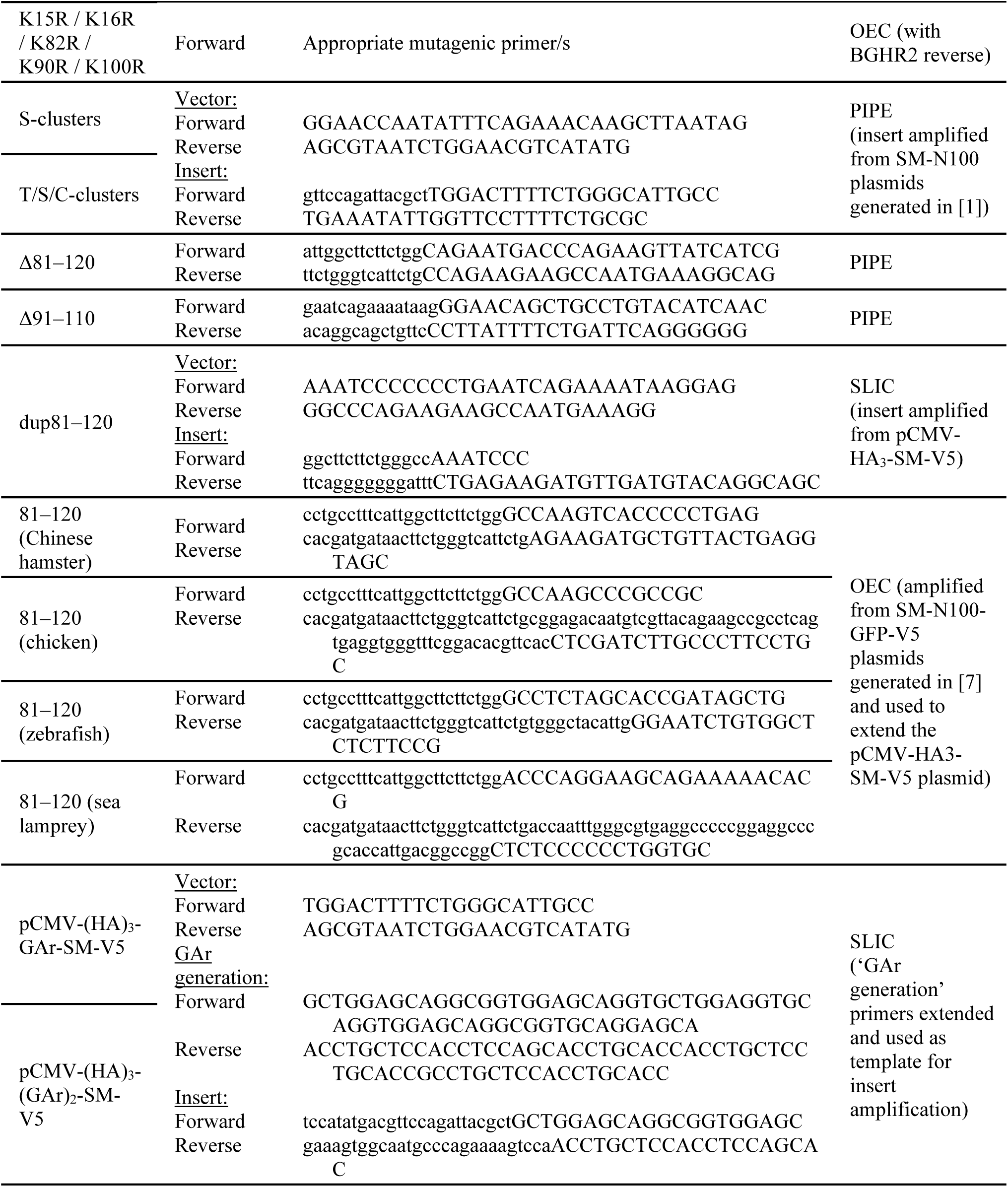
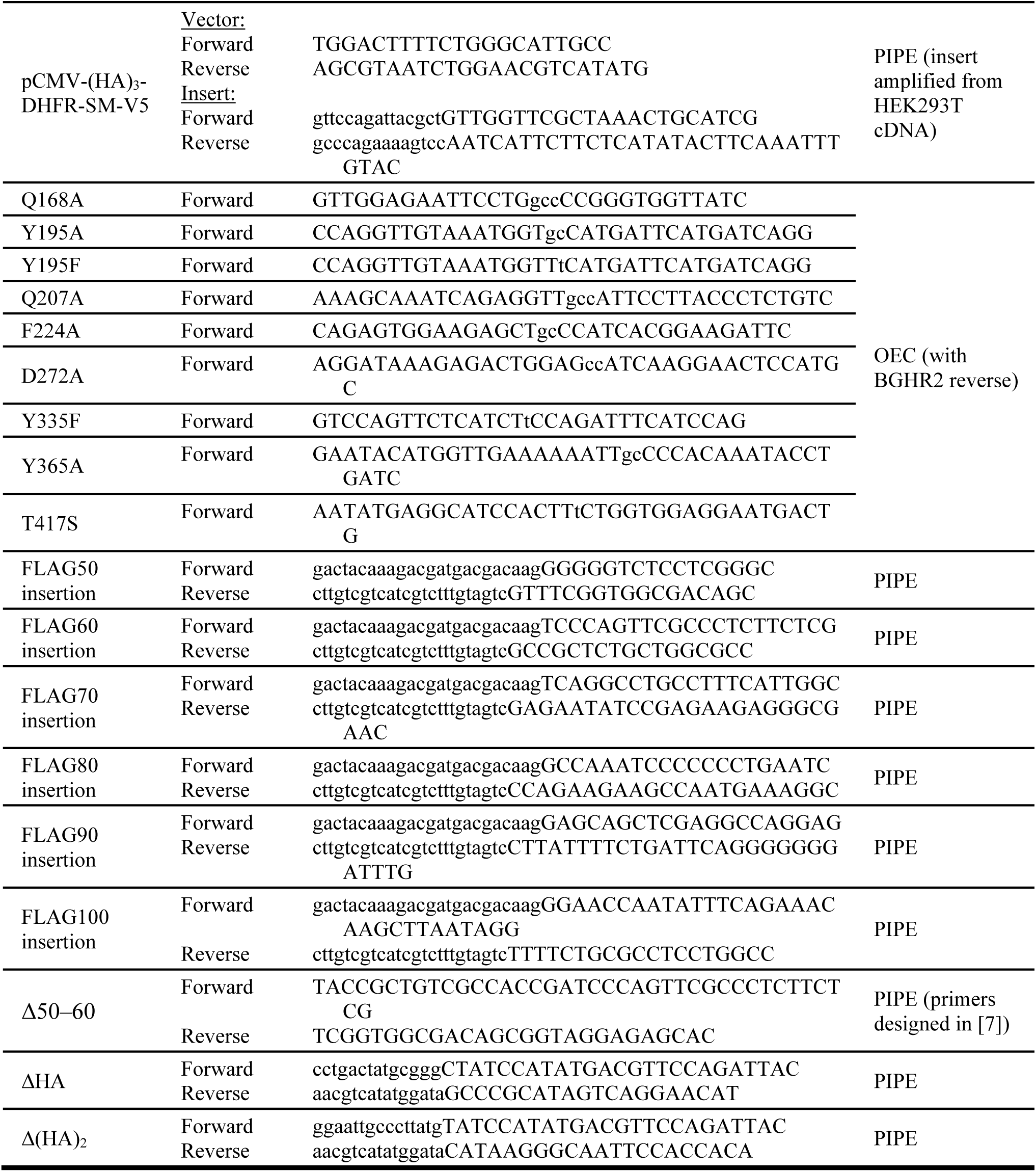
Primers used for qRT-PCR and DNA cloning. Non-annealing nucleotides for DNA insertions, deletions and substitutions are indicated in lowercase. Abbreviations for cloning methods: OEC (overlap extension cloning) [4] PIPE (polymerase incomplete primer extension cloning) [5]; SLIC (sequence- and ligation-independent cloning) [6].

## Notes

### Competing Interest Statement

The authors have declared no competing interest.

